# Leptin: A Potential Adipokine Bridging Systemic Endotoxemia and Sex-Divergent Temporomandibular Joint Osteoarthritis Development

**DOI:** 10.1101/2025.10.13.682179

**Authors:** Shipin Zhang, Siyi Chen, Marina Fonti, David Fercher

## Abstract

**Background:** Temporomandibular joint (TMJ) osteoarthritis (OA) is a degenerative disease affecting the whole synovial joint, with a higher prevalence in women. Obesity is recognised as a risk factor for knee OA, but its association with TMJ degeneration remains controversial. Lipopolysaccharide (LPS), or endotoxin, has increasingly been proposed as a mediator of obesity-related knee OA. In parallel, adipose-derived leptin played a role in the development of metabolic OA. This study investigated whether chronic LPS exposure induces TMJ OA and whether leptin is involved in this process.

**Methods:** Six-month-old female and male Wistar rats received continuous subcutaneous LPS infusion to induce TMJ OA. At 10 weeks, peripheral blood, white adipose tissue, and TMJs were harvested for biochemical, histological, micro-computed tomography and gene expression analyses. Primary TMJ chondrocytes isolated from healthy rats were used to investigate sex-specific cellular responses to leptin and LPS *in vitro*.

**Results:** Chronic LPS exposure induced mild, sex-divergent TMJ changes. Compared to same-sex healthy controls, LPS-treated females showed a marginally worse Mankin score (*P* = 0.053) with greater cartilage glycosaminoglycan loss (*P* = 0.032), synovitis and subchondral bone deterioration, whereas LPS-treated males only showed worse cartilage surface fibrillation. Both sexes showed subcutaneous adipocyte hypertrophy in response to LPS. Only females progressed to adipose inflammation and plasma leptin elevation (*P* = 0.018), which correlated strongly with total Mankin score (Spearman rho=+0.773, *P* = 0.024). Leptin was detected in the joint on chondrocytes co-expressing leptin receptor (OB-R) and inducible nitric oxide synthase (iNOS). The percentage of cells positive for leptin, OB-R and iNOS was differentially abundant in LPS-treated females than in sex-matched controls and LPS-treated males. *In vitro*, at the same leptin dose, female chondrocytes exhibited lower metabolic activity and greater oxidative stress than male chondrocytes. Notably, leptin upregulated matrix metallopeptidase (MMP)-13 expression only in LPS-primed female chondrocytes.

**Conclusion:** Chronic systemic LPS exposure induced sex-divergent TMJ osteoarthritic pathology, adipose dysfunction and altered leptin signalling. The tissue-level changes were associated with leptin-linked inflammatory cascades amplified in female chondrocytes. These findings suggest that leptin is involved in the pathogenesis of systemic endotoxin-triggered TMJ OA and may partly explain sex differences in OA development.

## 1. Introduction

The temporomandibular joint (TMJ) is a synovial joint in the craniofacial region that experiences mechanical loading when functioning (1,2). Osteoarthritis (OA) is a degenerative disease that can affect all synovial joints, including the TMJ. Pathological changes of OA include cartilage surface fibrillation, glycosaminoglycan (GAG) loss, bone remodelling and synovitis, and is significantly more prevalent in females than males (2–4).

Recently, metabolic osteoarthritis (MetOA) has been recognized as a subtype of OA that is associated with systemic low-grade inflammation, oxidative stress, adipose dysregulation, and other metabolic changes (5–7), often linked to high body mass index (BMI) and obesity (8,9). While the association between obesity and knee OA is well established, the relationship between body mass and TMJ OA remains controversial. Population studies have suggested that high BMI is not a risk factor for TMJ disorders (TMD) and may even be a protective factor for TMD pain (10–12), whereas preclinical results have shown that high-fat-diet (HFD)-induced obesity promotes TMJ OA (13,14). The discrepancy between human and animal studies suggests high body mass alone may not fully explain the relationship between obesity and TMJ OA. This raises the possibility that obesity-associated systemic factors, rather than increased body mass per se, drive the development of TMJ OA.

Metabolic endotoxemia, defined as a subclinical elevation of circulating endotoxin (lipopolysaccharide, LPS) (15), has emerged as a potential key mediator. Elevated LPS levels have been observed in obese individuals (16–18) and diet-induced obese rodents (15,19,20). LPS has been implicated in the pathogenesis of knee arthritis (15,18,21–23) by inducing systemic inflammation and modulating immune responses (24). In addition, LPS was reported to stimulate the release of leptin (25,26), a proinflammatory adipokine secreted primarily by subcutaneous adipose tissue rather than intra-abdominal adipose tissue (27,28). The level of circulating leptin is generally proportional to total fat mass and has a stronger correlation with adiposity in women than in men (27,29). Recent studies have shown that leptin was present in the synovial fluid of TMJ-OA patients (30), and was proven to mediate knee OA progression and pain (31). However, whether LPS contributes to TMJ OA and how it interacts with leptin in this context remains unclear. Furthermore, the mechanisms underlying sex differences in OA development are not fully understood.

To address these gaps, we investigated the effects of chronic systemic LPS exposure on TMJ OA development in male and female rats, and the involvement of adipose tissue and leptin in this process. We administered LPS systemically for 6 weeks and subsequently assessed osteoarthritic changes in the TMJ, systemic inflammatory markers, adipose tissue alterations, and adipokine profiles within each biological sex. Additionally, we evaluated sex-divergent effects of LPS and leptin on female and male TMJ condylar chondrocytes *in vitro*.

## 2. Method

The detailed study protocols were included in the Supplementary Materials. The Primers used in this study were listed in Supplementary Table 1.

### 2.1. Animal experiment

The animal study was approved by the Veterinary Office of the Canton Zürich (License No. ZH158/2021) and conducted in accordance with ARRIVE guidelines. Thirteen male and fourteen female Wistar rats (6 months old, Janvier Labs, France) with a mean weight of male:485 g (372 – 632 g) and female: 291 g (253 – 330 g) were used. A pilot trial involving 6 females and 5 males was conducted to determine the animals’ tolerance to chronic LPS infusion and to calculate the required sample size to achieve 80% study power. In this trial, animals were randomly allocated into the LPS group (2 males and 3 females) and the saline control group (3 males and 3 females). Rats were implanted subcutaneously with osmotic pumps (Alzet 2ML4, DURECT Corporate, USA), as previously described (32). The pump was either filled with 0.9% Sodium chloride (NaCl) (B.Braun, Germany) or LPS from Escherichia coli (*E. coli*) O26:B6 (Sigma-Aldrich, USA) to infuse 18 µg/kg/day LPS for 4 weeks (Figure 1A). In the main trial, the animals were randomised into Control or LPS group (n = 4 per sex and group) by block randomization method (33). Osmotic pumps (Alzet 2006) were used to continuously deliver either 0.9% NaCl or LPS at 18 µg/kg/day for 6 weeks (Figure 1B). At the end of LPS infusion, the osmotic pumps were left in place in the animals for 6 weeks (pilot trial) or 4 weeks (main trial) till the end of the experiment. In both trials, animals were housed in groups of 2 or 3 in individually ventilated cages in a standard laboratory animal environment (21± 3 °C and 12/12h light–dark cycle), received acidified tap water (pH 2.5-3.0) and standard laboratory rodent diet (KLIBA NAFAG, CH) in a specific opportunistic pathogen-free facility. After ten weeks, animals were euthanized by CO_2_ overdose. Whole blood, bilateral TMJ condyles, rat heads, dorsocervical subcutaneous adipose tissue (SAT) and the mesenteric visceral adipose tissue (VAT) were collected (34). The right TMJ condyles collected from the pilot trial, intended for micro-computed tomography (µCT) and histological analysis, were excluded from the study due to a sample-preparation error. The left TMJ condyles from the pilot trial were subjected to gene expression analysis. Two blood samples were excluded from LPS measurement due to haemolysis. All the other tissues were included in their respective analyses, as detailed in Figure 1 and the method sections below. For the *in vitro* mechanistic study, primary TMJ condylar chondrocytes were isolated from 8-month-old healthy rat carcasses (2 males and 2 females) obtained as surplus material from another study (Figure 1C).

**Figure 1.**
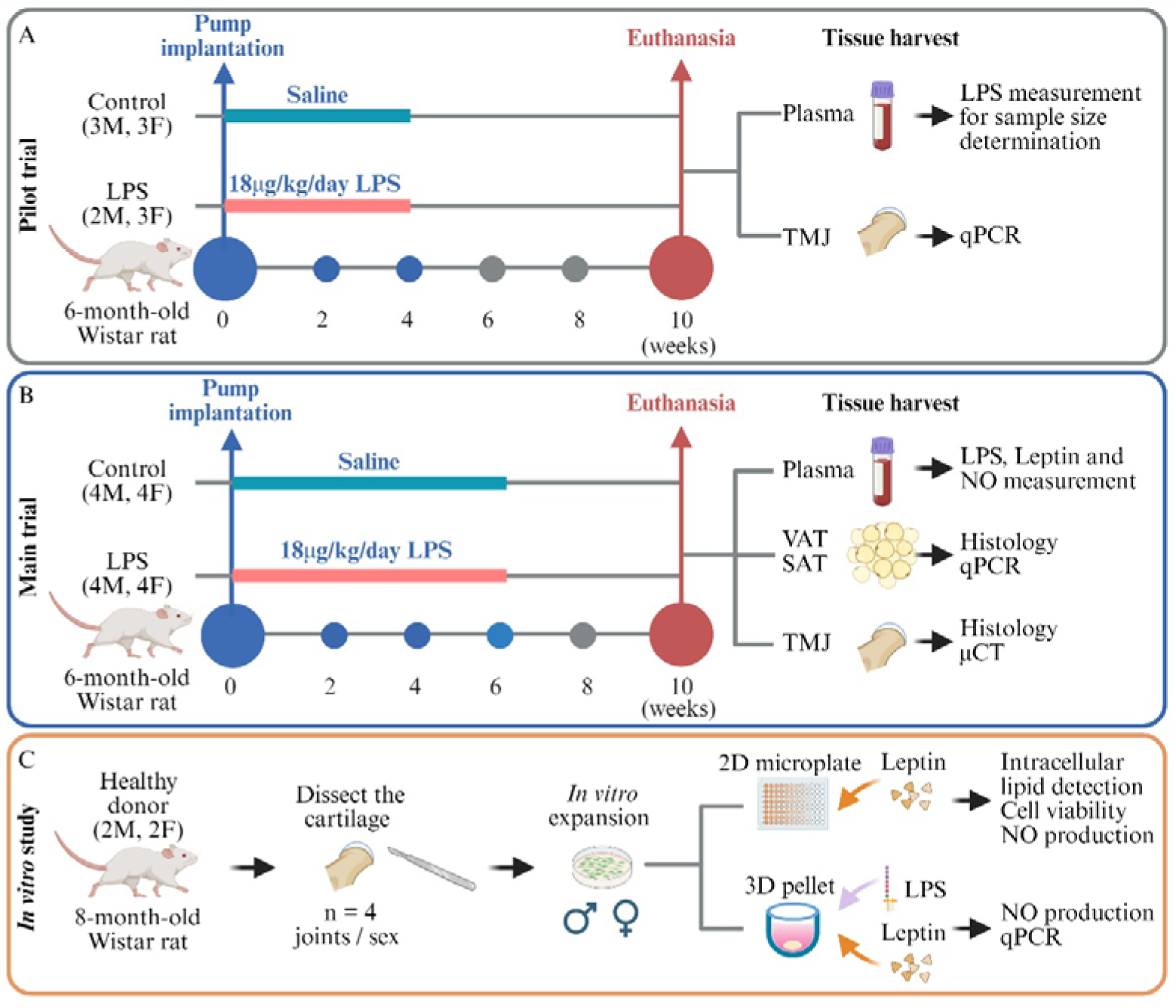
Schematic illustration of in vivo and in vitro experiment design. Schematic of animal groups, study timeline and tissue harvest plans for (A) pilot trial and (B) main trial. (C) *In vitro* experimental setup to study the role of leptin in OA development. M: Male; F: Female. In the pilot trial, blood was collected to measure LPS concentration, and TMJ condyles were harvested for qPCR-based gene expression analysis. In the main trial, blood was collected to measure LPS, nitric oxide (NO), and leptin. Subcutaneous adipose tissue (SAT) and visceral adipose tissue (VAT) were collected for qPCR-based gene expression analysis and histomorphometry evaluation. The heads were collected for micro-computed tomography bone analysis and histology evaluation. For the *in vitro* study, TMJ condylar chondrocytes were isolated from healthy rat joints and used for mechanistic evaluations. Created in BioRender. Zhang, S. (2026) https://BioRender.com/vdjaeod

### 2.2. Blood analysis

5 mL whole blood was collected post-mortem via cardiac puncture into a BD Vacutainer® EDTA Tubes (BD, USA), centrifuged for 10 min at 1500 g to remove cells, then for 10 min at 2000 g to deplete platelets from plasma. Plasma was diluted 50-fold with endotoxin-free water for LPS concentration analysis by the LAL assay kit (Thermofisher), 30-fold for nitric oxide (NO) quantification using Griess reagent kit (Thermofisher), or 3-fold for leptin quantification using an ELISA kit (Merck). Assays were performed according to manufacturers’ instructions.

### 2.3. Micro-computed tomography evaluation

The heads were dissected and fixed at 4% (v/v) paraformaldehyde (PFA). High-resolution µCT scan (µCT45, SCANCO MEDICAL) was performed to assess bone morphology. For bone parameter analysis, a stack of 200 coronal slides across the entire condyle were selected. The percentage of bone volume over total volume (BV/TV, %), trabecular thickness (Tb.Th, µm), trabecular separation (Tb.Sp, µm), trabecular number (Tb.N, 1/mm), and bone mineral density (BMD) were quantified using CTAn version 1.13 (Bruker microCT, Kontich, Belgium).

### 2.4. Histomorphometric analysis

SAT, VAT and TMJs were paraffin-embedded (35), cut at 5 µm, and stained with haematoxylin and eosin (HE) for general morphology. TMJs were stained for Safranin-O/fast green (Saf-O) for GAG deposition, picrosirius red (PSR) for fiber orientation (36), and immunohistochemistry (IHC) to detect the distribution of type II collagen (Col II) (37) or inducible nitric oxide (iNOS) in condylar cartilage (38). Immunofluorescent (IF) staining was performed to detect the expression and colocalization of iNOS, leptin and leptin receptor (OB-R) in chondrocytes. Mankin scoring (39) was performed on Saf-O-stained TMJs to evaluate cartilage degeneration. Krenn’s synovitis score (40) was performed on HE-stained slides to assess the synovial inflammation. Both scorings were performed independently by two blinded observers (S.Z and M.F). QuPath v0 and ImageJ software (National Institutes of Health) were used for semi-quantitative image analysis of adipocyte size, Saf-O- and Col II-stained area, percentage of iNOS-positive (iNOS^+^), Leptin-positive cells (Leptin^+^), OB-R-positive cells (OB-R^+^), and iNOS/Leptin/OB-R triple-positive (Triple^+^) cells over total cells. The collagen fiber orientation in condylar cartilage was measured using OrientationJ (41).

### 2.5. *In vitro* mechanistic study

TMJ condylar chondrocytes were isolated and cultured using a method adapted from Zhang et al (38). For the 2D microplate assay, 5000 chondrocytes were seeded per well in a 96-well plate overnight to allow cell adhesion, then stimulated with 1, 2, 5, 10, or 20 ng/ml of rat leptin (PeproTech) for 24 hours. At the end of the treatment, MTS assay (Abcam) was performed to measure the cellular metabolism. Cell culture supernatant was collected for NO quantification. Nile red staining was performed on the cells to visualise intracellular lipid droplets. For the 3D pellet assay, the chondrocyte pellets were prepared (38), and stimulated with 100 ng/ml LPS for 24 hours, followed by 20 ng/ml leptin for 24 hours. The culture supernatant was collected for NO quantification. The chondrocyte pellets were lysed for Picogreen DNA quantification (Thermofisher) per the manufacturer’s instructions and for gene expression analysis (Figure 1C).

### 2.6. RNA isolation and qRT-PCR analysis

RNA extraction from chondrocyte pellets was performed with the RNeasy^®^ kit (Qiagen) according to the manufacturer’s instructions. RNA extraction from adipose tissue and TMJ condyles was performed as previously described ^29^. Gene expression heatmap of TMJ condyles was generated using Morpheus (https://software.broadinstitute.org/morpheus/). Samples were hierarchically clustered using one minus Pearson correlation as the metric and average linkage as the linkage method, and stratified by sex. Genomic DNA was extracted from chondrocytes to verify the biological sex of cells using a single-step PCR-based method (42).

### 2.7. Statistics

Blood LPS concentration measured from the pilot trial (Supplementary Table 2) was used to calculate the sample size using G*Power 3.1.3 (43,44). A minimal of three animals per group per sex were estimated to provide 80% power with a significance level of *p* = 0.05 (two-tailed) to detect differences in circulating endotoxin between treatments in females and between sexes in the LPS group. Therefore, four animals per group per sex were included in the main trial. Each animal is counted as one experimental unit for blood analysis, adipose size quantification, and weight change. Left and right joints from the same animal were averaged and treated as a single biological replicate for semi-quantification of Saf-O^+^ area, Col II^+^ area, % iNOS^+^ in the whole cartilage, histological scorings, fiber orientation analysis, µCT evaluation and IF quantification. Unless otherwise stated, all data were presented as mean (95% confidence interval (CI)). For continuous outcomes, normality was assessed per group using Shapiro-Wilk test. For data that met the assumption of normality, Levene’s test was used to evaluate the equality of variances. Student’s t-test was applied where variances were equal, and Welch’s t-test where variances were unequal. For ordinal outcomes or data that failed normality testing, the Mann-Whitney U test was used for pairwise comparisons; the Scheirer-Ray-Hare test was used for non-parametric factorial comparisons. Effect sizes are reported as Hedges’ g for parametric comparisons or the Hodges-Lehmann location shift estimate (HL) and rank-biserial correlation (r_rb) for non-parametric comparisons. Exact 95% confidence intervals for HL were derived from the Wilcoxon rank-sum critical value (k=5 for n =n =4); 95% CIs for r_rb were computed via Fisher’s z-transformation. Prior to the assessment of the main effects of Sex, Treatment and Sex × Treatment interaction on the outcomes, ANOVA model residuals were subjected to Shapiro-Wilk normality test to check the data distribution. For in vivo gene expression data (fold-change relative to Female SAT control), values were log2-transformed prior to normality test and subsequent analysis. A Two-way ANOVA (Sex × Treatment) was fitted for outcomes that passed the Shapiro-Wilk normality test. Type II sums of squares were used, yielding F-statistics and *P* value with df = (1,12). Effect sizes were expressed as partial η². For ordinal outcomes (Mankin scores and Krenn’s synovitis score) or continuous outcome that failed the normality assumption, the Scheirer-Ray-Hare test was applied. For each effect, the H statistic was computed and evaluated against a chi-square distribution with df = 1. For the forest plot effect sizes, bias-corrected and accelerated bootstrap confidence intervals were computed using 2000 stratified resamples (45). For correlation analysis, Spearman’s rank correlation coefficient (rho) was used. The interclass correlation coefficient (ICC) was calculated to assess the agreement between the two independent observers’ histopathological scores. The statistical significance level was set at *P* < 0.05. SPSS version 29.0 (IBM, Chicago, IL, USA) was used for ICC calculation. Analysis of the data generated from TMJ tissue gene expression and *in vitro* experiment were performed in GraphPad Prism 10 (GraphPad Software, LLC). Python (Version 3.12) was used for all the other data analysis.

## 3. Results

### 3.1. LPS infusion induced sex-dimorphic degenerative changes in TMJ cartilage, bone and synovium

#### 3.1.1. Cartilage

We observed mild, sex-dimorphic osteoarthritic changes in the TMJ condylar cartilage of LPS-treated. LPS-infused male animals showed slight surface fibrillation and disruption of cartilage surface structure (Figure 2A and Supplementary Figure 1). In comparison, LPS-treated females appeared to have a greater loss of cartilage matrix throughout the cartilage layer (Figure 2A and Supplementary Figure 2). Mankin scores of condylar cartilages first indicated that female rats without LPS treatment already trended towards worse total Mankin scores than male controls (HL = 1.38, 95%CI: 0.50 to 1.75, r_rb: 0.75, *P* = 0.110), suggesting a higher baseline vulnerability in healthy musculoskeletal mature females than males. LPS treatment resulted in greater absolute increment in Mankin scores than sex-matched controls in males (HL: 2.25, 95% CI: 1.50 to 3.50, *P* = 0.108) than in females (HL: 1.00, 95% CI: 0.75 to 1.25, *P* = 0.053), demonstrating a significant catabolic effect on cartilage integrity (H (1,12) = 7.52, *P* = 0.006) in both sexes (Figure 2B). Comparing between females and males, the effect of LPS on female rats were smaller but more consistent than that on males. All four LPS-treated females score at or above the highest female control value. Whereas, three of four LPS-treated males exceeded male control values, with one overlapping the lower control range. This observation was supported by the higher rank-biserial correlation effect size of LPS in females (r_rb: 0.88, 95% CI: 0.44 to 0.98) compared to males (r_rb: 0.75, 95% CI: 0.10 to 0.95). The increase in the total Mankin score was attributed to changes in different subcategories in each sex. LPS-treated males showed marginally worse structural regularity (HL: 0.50, 95% CI: 0.25 to 1.00, *P* = 0.055) (Supplementary Figure 1B), whereas LPS-treated females showed a trend toward reduced cartilage matrix staining (HL: 0.75, 95% CI: 0.25 to 1.00, *P* = 0.080) (Supplementary Figure 2B).

**Figure 2.**
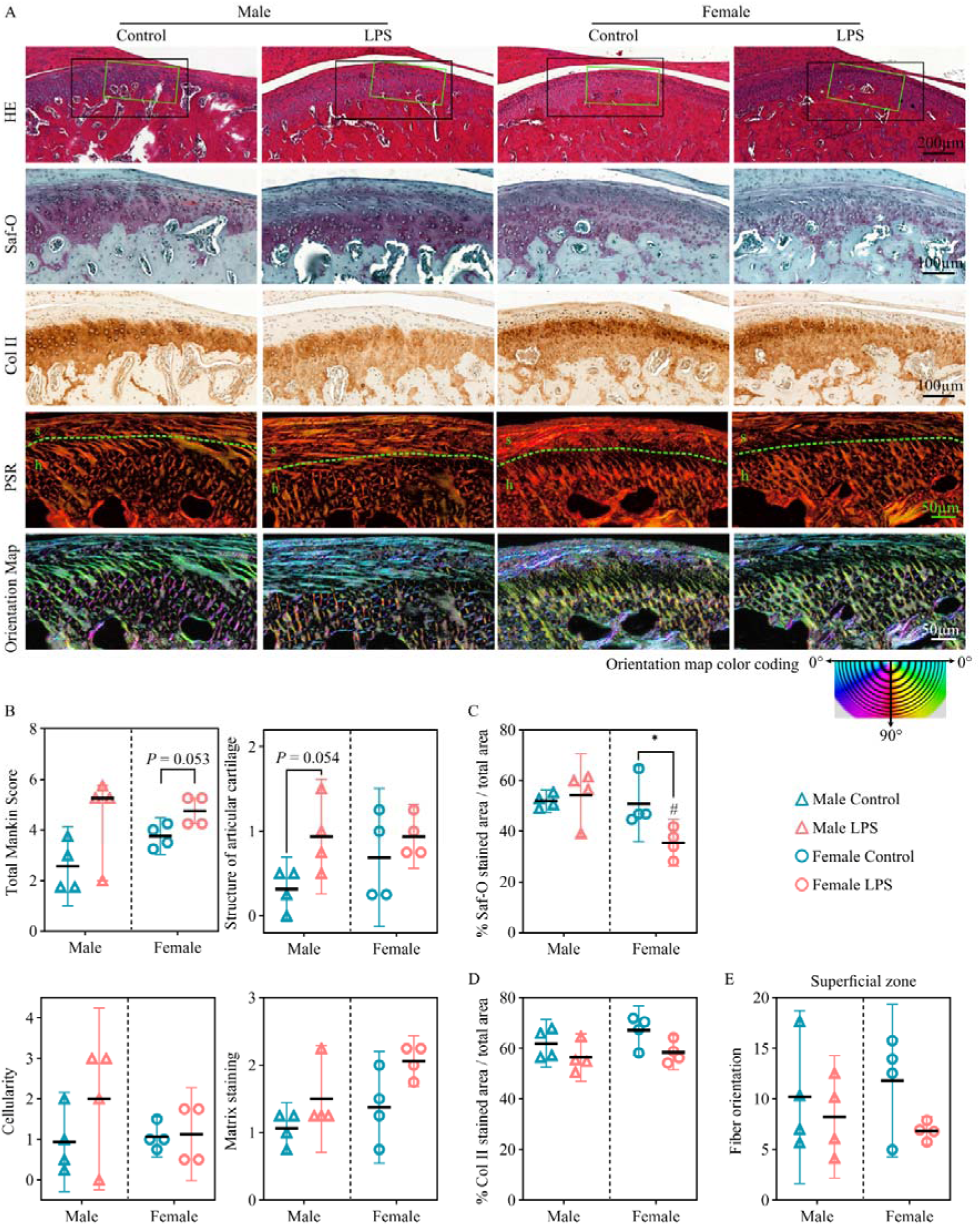
LPS infusion is associated with sex-dimorphic degenerative changes in TMJ condylar cartilage. (A) Representative images of TMJ stained with hematoxylin and eosin (HE), safranin-O and fast green (Saf-O), collagen II (Col II), picrosirius red (PSR) imaged using a polarized microscope with corresponding pseudo-colour orientation maps of PSR to visualise the fibre orientation (s: superficial zone; h: hypertrophic zone). Black boxes in the HE image indicate the regions where high magnification Saf-O and Col II images were taken, whereas the green boxes indicate the region where high magnification PSR images were taken. Scale bar: 200µm (HE), 100 µm (Saf-O and Col II), 50 µm (PSR and orientation map). (B) Total Mankin score and subcategory scores of the structure of articular cartilage, cellularity and matrix staining. The total Mankin scores are presented as median ± 95%CI with individual data points. Inter-rater reliability between the two independent observers was good (ICC = 0.83, 95%CI: 0.60 – 0.92, *P* < 0.001). The Mankin subcategory scores are presented as mean ± 95%CI with individual data points. Comparisons performed using Mann-Whitney U tests; Hodges-Lehmann (HL) location shift estimates with 95%CI, r_rb and exact P values are reported in the text. * *P* < 0.05 compared to the control group within the same sex. N = 4 animals per group and sex. Quantification of (C) safranin-O-stained area (Saf-O^+^) and (D) collagen II-stained area (Col II^+^) in condylar cartilage. (E) Fibre orientation of the superficial zone. Data in panels C–E are presented as mean ± 95% CI, with individual data points. Comparisons performed using Student’s t test; mean difference with 95%CI, Hedges’ g, and exact P values are reported in the text. * *P* < 0.05 compared to the control group within the same sex. N = 4 animals per group and sex.

The semi-quantitative Mankin score was further supported by areal quantification of GAG and collagen II deposition in condylar cartilage. Relative to same-sex controls, LPS-treated female rats had significantly less Saf-O^+^ area (mean difference: -15.34%, 95%CI: -28.86 – - 1.81%, Hedges’ g = -1.71, *P* = 0.032), and showed a moderate reduction in Col II^+^ area (mean difference: -8.63%, 95%CI: −17.89 to 0.62%, Hedges’ g = -1.40, *P* = 0.063) in the cartilage. In males, LPS-associated changes in the cartilage ECM varied greatly among individuals and the trends were inconclusive. Compared to controls, LPS-treated males exhibited slightly more Saf-O^+^ (mean difference: 2.25%, 95%CI: -10.80 to 15.29%, Hedges’ g = 0.259, *P* = 0.689) but less Col II^+^ (mean difference: -5.52%, 95%CI: -15.79 to 4.75%, Hedges’ g = -0.81, *P* = 0.236) (Figure 2C, D). Two-way ANOVA analysis suggested that the changes in % Saf-O^+^ was significantly influenced by the combined effect of Sex and Treatment (F (1,12) = 5.24, partial η²: 0.30, *P* = 0.041), confirming the effect of LPS on GAG loss differ by sex. Whereas LPS treatment has a dominant effect on % Col II^+^ dominantly (F(1,12) = 6.28, partial η²: 0.34, *P* = 0.028) over Sex (F(1,12) = 1.60, partial η²: 0.12, *P* = 0.230) and Sex × Treatment interaction (F (1,12) = 0.30, partial η²: 0.025, *P* = 0.592), supporting our finding showing LPS uniformly reduce the collagen II content in male and female cartilage. Collagen fibre derangement was visualised in pseudo-colour maps based on polarized PSR images (Figure 2A and Supplementary Figure 3). On average, superficial zone collagen fibres in LPS-treated females displayed a lower orientation angle relative to the cartilage surface than those in control females (6.84° vs 11.81°, mean difference: -4.97°, 95% CI: -10.87 to 0.93°, Hedges’ g: -1.27, *P* = 0.085). A smaller change in fiber orientation was observed in males (mean difference: -1.94°, 95% CI: -10.00 to 6.11°, Hedges’ g: -0.36, *P* = 0.576) (Figure 2E).

#### 3.1.2. Subchondral bone

The TMJ condylar head bone structure was visualised and quantified by µCT (Figure 3A). Significant main effects of biological sex were detected for BV/TV (F(1,12) = 47.5, *P* < 0.001, partial η² = 0.798), trabecular thickness (Tb.Th: F(1,12) = 75.44, *P* < 0.001, partial η² = 0.863), and trabecular separation (Tb.Sp: F(1,12) = 39.34, *P* < 0.001, partial η² = 0.766), explaining a larger proportion of variance than LPS treatment in all three parameters across all animals (partial η²=0.076 to 0.208; Figure 3B-D). Despite the dominant effect of biological sex, LPS exposure was associated with female-specific deterioration of trabecular microstructures, with no corresponding changes detected in males. Relative to female controls, LPS-treated females showed a significant 11% increase in trabecular separation (Tb.Sp: mean difference 0.0041 mm, 95%CI: 0.0002 to 0.0081, Hedges’ g = 1.88, *P* = 0.045) (Figure 3D) and a significant 2% reduction in trabecular number (Tb.N: mean difference: -0.22/mm, 95%CI: -0.41 to -0.03, Hedges’ g = -1.73, *P* = 0.031) (Figure 3E). Neither of these changes was significant in males. BV/TV trended lower in both sexes following LPS infusion but did not reach statistical significance (Male: -1.89%, 95%CI: -5.51 to 1.73%, Hedges’ g = -0.79, *P* = 0.248; Female: -1.13%, 95%CI: -3.91 to 0.93%, Hedges’ g = -0.83, *P* = 0.228) (Figure 3B). No difference was detected in bone mineral density (BMD) between sexes or across groups (Figure 3F).

**Figure 3.**
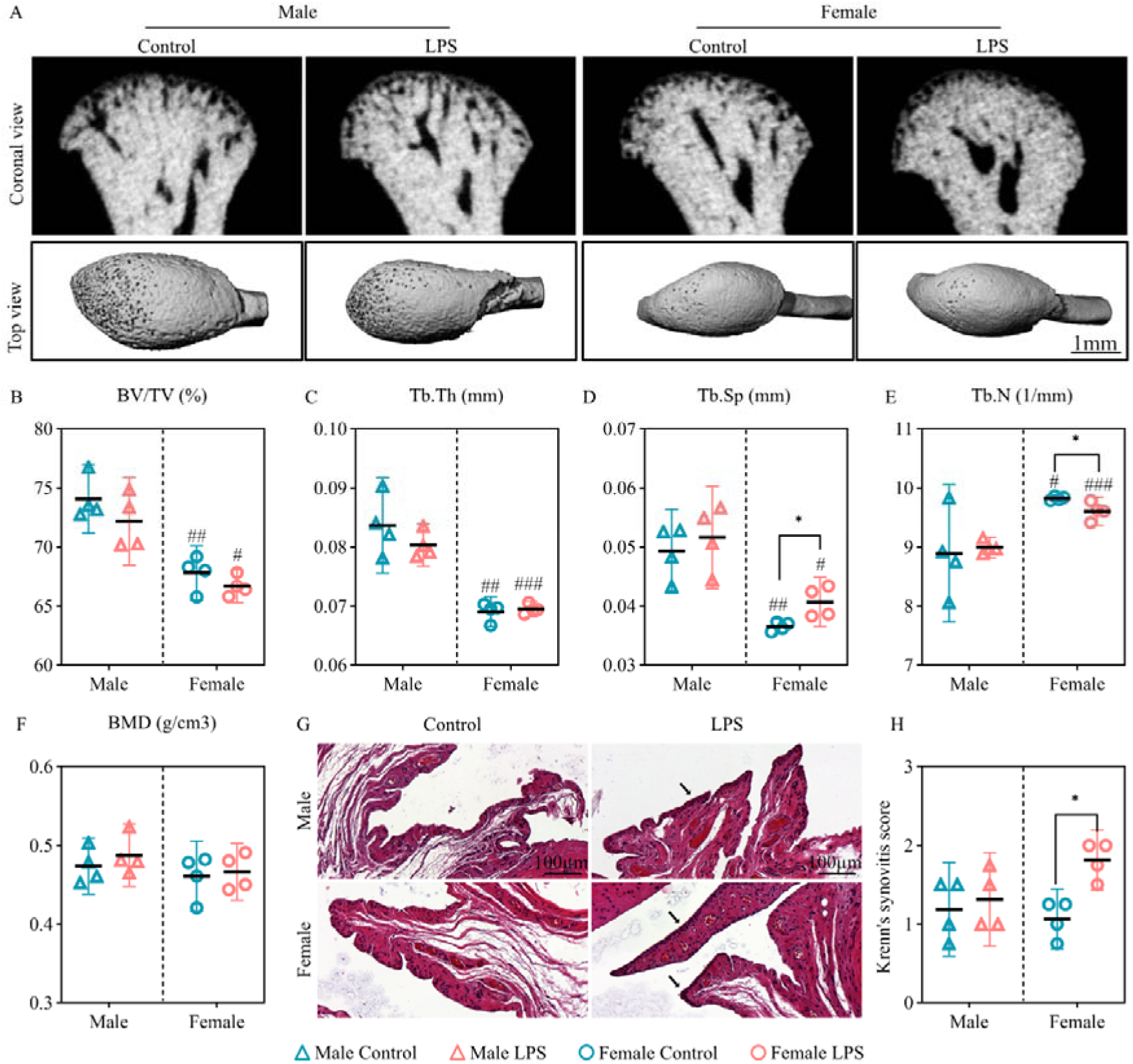
Assessment of subchondral bone microstructure and synovial membrane inflammation in TMJ. (A) Representative µCT images of TMJ condyles in coronal and top views. Scale bar: 1 mm. Quantitative analysis of bone parameters: (B) Percentage of bone volume over total volume (% BV/TV); (C) Trabecular bone thickness (Tb.Th); (D) Trabecular bone separation (Tb.Sp); (E) Trabecular bone number (Tb.N) and (F) Bone mineral density (BMD). Comparisons performed using Student’s t test; mean difference with 95%CI, Hedges’ g, and exact P values are reported in the text. (G) Histological analysis of TMJ synovial membrane using HE staining; Black arrow indicates the increase of the synovial lining layer. Scale bar: 100 µm. (H) Krenn’s synovitis score used to assess overall synovial inflammation. N = 4 rats per group and sex. Data represent mean with 95% CI. Comparisons performed using Mann-Whitney U tests; Hodges-Lehmann (HL) location shift estimates with 95%CI, r_rb and exact P values are reported in the text. For all graphs, data represent mean with 95% CI, with each individual data point shown (n = 4 per group per sex). * *P* < 0.05 compared to control group within same sex. ^#^ *P* < 0.05, ^##^ *P* < 0.01, ^###^ *P* < 0.001 compared to the males within the same treatment group.

#### 3.1.3. Synovium

Synovial inflammation was assessed on HE-stained synovial membrane (Figure 3G and Supplementary Figure 4) using Krenn’s synovitis scoring system (Figure 3H and Supplementary Figure 4C,D). Systemic LPS exposure generally elevated synovial inflammation across all animals (Scheirer-Ray-Hare H = 4.88, *P* = 0.027), with the response markedly greater in females. LPS-treated females showed a significantly higher total Krenn’s score than female controls (HL: 0.75, 95% CI: 0.50 to 1.00, r_rb = 1.00, *P* = 0.028). In contrast, LPS infusion did not consistently increase synovitis scores in male rats (HL = 0.125, 95% CI: 0.00 to 0.25, r_rb = 0.25, *P* = 0.649) (Figure 3H).

To assess the effects of LPS on three major OA joint pathologies measured by Mankin score and cartilage matrix staining, Krenn’s synovitis score and subchondral bone parameters, we performed Cohen’s d effect size estimation within each biological sex (Control and LPS pooled standard deviation, df = 6). LPS perturbation was estimated to have significant large effects in female compared to female controls in Mankin score (Cohen’s d: +1.92, 95%CI: +0.60 to +3.00, *P* = 0.037), % Saf-O^+^ area (Cohen’s d: -1.96, 95%CI: -3.63 to -1.01, *P* = 0.039), total Krenn’s score (Cohen’s d: +3.13, 95%CI: +1.57 to +4.18, *P* = 0.004), and Tb.Sp (Cohen’s d: +2.16, 95%CI: +0.96 to 3.40, *P* = 0.045). Similar large effect of LPS on Mankin score was observed in males but did not reach a statistically significance level (Cohen’s d: +1.42, 95%CI: -0.27 to +2.27, *P* = 0.103) (Supplementary Figure 5). In brief, continuous LPS infusion induced mild TMJ degenerative changes, affecting cartilage surface integrity, ECM content, synovium homeostasis, and subchondral bone microarchitecture in a sex-dependent manner, with greater consistency in female rats.

### 3.2. LPS infusion induced sex-divergent adipocyte hypertrophy, adipose inflammation and systemic leptin and NO elevation

Having established that LPS infusion induced sex-divergent joint pathology, we next examined whether there were differences in systemic and adipose tissue responses to LPS, which could potentially account for this divergence. At the study endpoint, 4 weeks after cessation of LPS infusion, plasma LPS concentration remained approximately 4-fold higher in LPS-treated female rats than in female controls, although this did not reach a statistically significant level (*P* = 0.080). No difference in plasma LPS concentration was observed in LPS-treated male rats compared to controls (Figure 4A). Despite the absence of a significant elevation in circulating LPS in either sex, LPS-treated rats showed an overall increasing trend in plasma NO concentration compared to controls (F(1,12) = 3.69, *P* = 0.079, partial η² = 0.235), suggesting that LPS infusion triggered a low-grade systemic inflammation. This effect reached statistical significance within females (mean difference: 20.83 µM, 95%CI: 9.85 to 31.81 µM, Hedges’ g = 2.85, *P* = 0.004) but not within males (mean difference: 6.67 µM, 95%CI: -26.59 to 39.92 µM, Hedges’ g = 0.30, *P* = 0.641) (Figure 4B).

**Figure 4.**
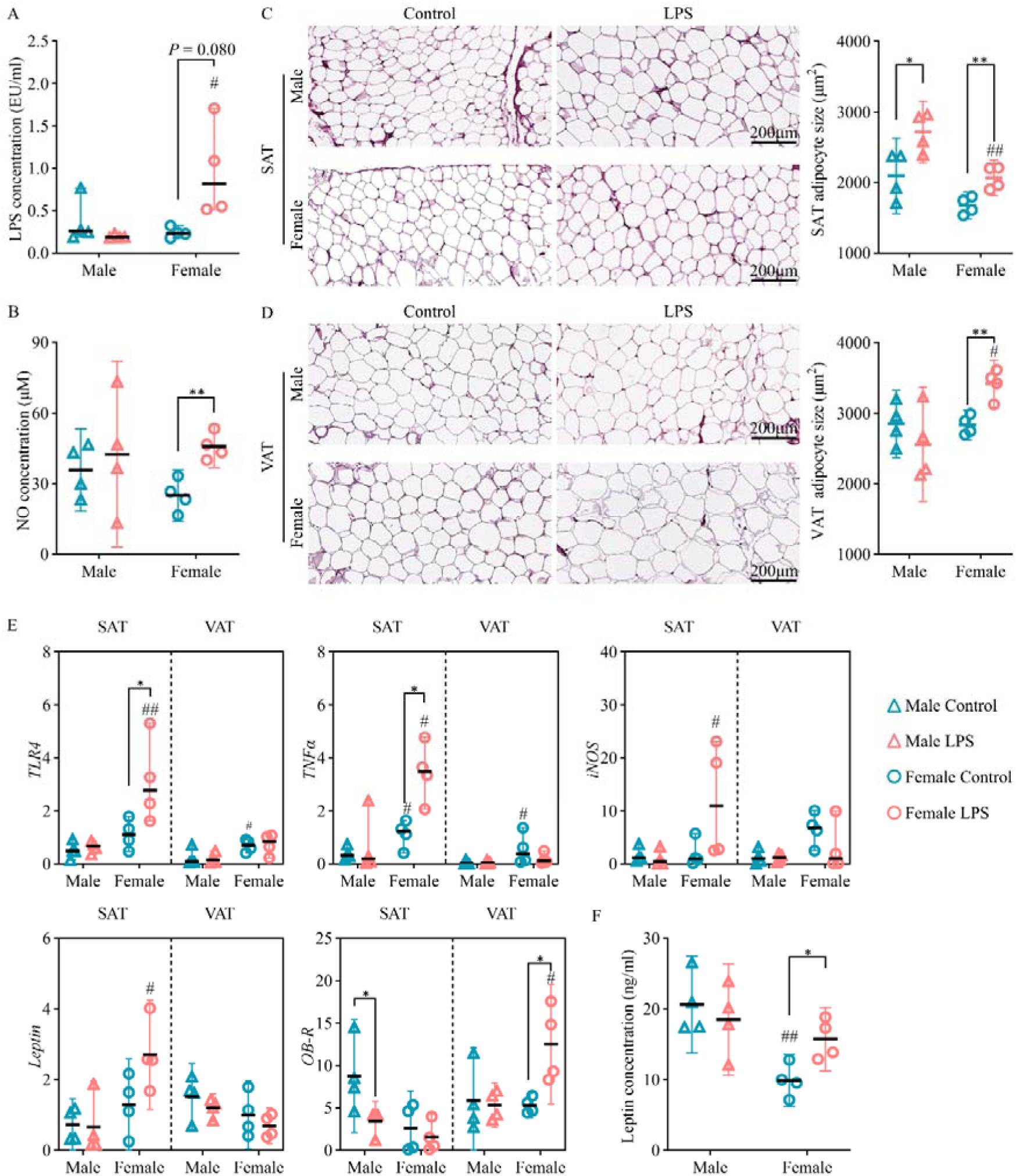
Sex differences in systemic and adipose tissue responses to LPS infusion. (A) Concentrations of LPS and (B) nitric oxide (NO) in blood plasma measured at the study endpoint. For plasma LPS, comparisons between female LPS and female control groups were performed using Welch’s t-test; Comparisons involving male groups and between sex comparisons within each treatment groups were performed using Mann-Whitney U test. For plasma NO, comparisons were performed using Student’s t-test. Representative haematoxylin and eosin (HE) staining and quantification of adipocyte size in (C) subcutaneous adipose tissue (SAT) and (D) visceral adipose tissue (VAT). Scale bar: 200 µm. Comparisons between treatment groups within each sex were performed using student’s t-test. Between-sex comparisons within each treatment group were performed using Welch’s t-test where indicating unequal variance (SAT, control groups), and Student’s t-tests otherwise. (E) Gene expression of *TLR4*, *TNF*_α_, *iNOS*, *Leptin* and *OB-R* in SAT and VAT, normalised against SAT female control. Gene expression fold-changes were log2-transformed prior to analysis to check the data distribution. For dataset that were normally distributed, student’s t-test (equal variances) or Welch’s t-test (unequal variances) was applied. *OB-R* in SAT for male LPS group failed normality test, all pairwise comparison involving this dataset was performed using Mann-Whitney U test. (F) Concentration of plasma leptin measured at the study endpoint. Comparisons were performed using Student’s t-test. For all graphs, data represent mean with 95% CI, with each individual data point shown (n = 4 per group per sex). * *P* < 0.05, ** *P* < 0.01 compared to the sex-matched control group. # *P* < 0.05, ## *P* < 0.01 compared to males within the same treatment group.

The development of low-grade inflammation in response to LPS infusion was accompanied by aberrant alterations in adipose tissue. Despite comparable body mass across groups at all measurement time points (Supplementary Figure 6), adipocyte hypertrophy in response to LPS occurred in a sex- and fat depot-dependent pattern. SAT adipocytes were significantly enlarged following LPS infusion in both male (mean difference: 621.50 µm^2^, 95%CI: 92.49 – 1150.51 µm^2^, Hedges’ g = 1.77, *P* = 0.03) and female rats (mean difference: 393.25 µm^2^, 95%CI: 148.14 – 638.36 µm^2^, Hedges’ g = 2.41, *P* = 0.008) (Figure 4C). In contrast, VAT adipocyte hypertrophy was observed in LPS-treated females (mean difference: 584.00 µm^2^, 95% CI: 280.00–887.00 µm^2^, Hedges’ g = 2.89, *P* = 0.003) and not in males (*P* = 0.360) (Figure 4D).

Sex differences were also evident in the adipose gene expression changes in response to LPS. Two-way ANOVA revealed a significant female-dominant sex effect on gene expression across four out of five genes examined in SAT: *TLR4* (partial η²: 0.59, *P* = 0.001), *OB-R* (partial η²: 0.38, *P* = 0.019), *Leptin* (partial η²: 0.40, *P* = 0.015), and *TNF*_α_ (partial η²: 0.55, *P* = 0.003). In female SAT, LPS infusion significantly upregulated LPS receptor *TLR4* expression approximately 3-fold compared to female controls (*P* = 0.035), suggesting a heightening in LPS sensitivity. In parallel with *TLR4* upregulation, the expression of proinflammatory marker *TNF*_α_ was significantly upregulated by 3-fold (*P* = 0.015) and *iNOS* showed a 7-fold elevating trend (*P* = 0.072) in LPS-treated females relative to control females. The expression of *Leptin* in female SAT also showed an increasing trend in LPS group compared to controls but did not reach a statistically significant level, while leptin receptor *OB-R* expression was similar between LPS-treated and control female (Figure 4E). In contrast, male SAT showed no change in the expression of *TLR4*, *TNF*_α_, *iNOS* and *Leptin* following LPS infusion. The only significant change in gene expression was observed for *OB-R*, which was significantly downregulated in the LPS group (3.43, 95%CI: 1.07 to 5.80) relative to male controls (8.76, 95%CI: 2.11 to 15.41, *P* = 0.029), reflecting a potential attenuation of leptin sensitivity in response to LPS infusion in male subcutaneous adipose tissue (Figure 4E). Comparing females and males within LPS group, female SAT showed markedly greater expression of *TLR4* (*>*4-fold, *P* = 0.003), *TNF*_α_ (14-fold, *P* = 0.024), *iNOS* (>24-fold, *P* = 0.036), and *Leptin* (∼7-fold, *P* = 0.020) than LPS-treated males, indicating a substantially greater adipose transcriptional response to LPS in females. A similar female-dominant sex main effect was observed in VAT for *TLR4* (partial η²: 0.51, *P* = 0.004), *TNF*_α_ (partial η²: 0.47, *P* = 0.007), *Leptin* (partial η²: 0.30, *P* = 0.040), and *OB-R* (partial η²: 0.29, *P* = 0.048), though the contrast between LPS and control groups within each sex were less pronounced than that in SAT.

The sex-divergent adipose responses extended beyond phenotypic remodelling and transcriptional activation, translating into persistent systemic increases in circulating leptin. Relative to female controls, the LPS-treated females showed a 60% increase in plasma leptin (mean difference: 5.88 ng/ml, 95% CI: 1.42–10.35 ng/ml, Hedges’ g = 1.98, *P* = 0.018). In contrast, LPS-treated males showed no significant difference relative to controls in plasma leptin (mean difference: -2.11 ng/ml, 95% CI: -10.16 to 5.93 ng/ml, Hedges’ g = -0.40, *P* = 0.544) (Figure 4F).

Collectively, these data show that systemic LPS infusion induced a low-grade systemic inflammation and lasting morphological change in adipose tissues in both sexes, confirming biological LPS exposure in males and females, but with different magnitude of effect. In females, this progressed to a coordinated response comprising transcriptional upregulation of genes encoding inflammatory cytokines and leptin in SAT, and sustained elevation of circulating leptin at the study endpoint. In males, SAT hypertrophy occurred alongside more limited transcriptional signature, with not corresponding change in circulating leptin.

### 3.3. Leptin was detected in TMJ cartilage and co-localized with OB-R- and iNOS-positive chondrocytes

Given the sex-specific elevation in circulating leptin observed in LPS-treated females, we next investigate whether leptin and its receptor OB-R were present within TMJ joint microenvironment, and whether their intra-articular distribution relates to the patterns of oxidative stress and cartilage damage. To answer these questions, we used iNOS as the oxidative stress marker known to contribute to OA pathogenesis (46). IHC and multi-color IF staining were conducted to characterize the expression of iNOS, leptin, and OB-R on condylar chondrocytes.

The iNOS^+^ chondrocytes were mainly located at the hypertrophic zones of the central and posterior regions of the TMJ condylar cartilage (Figure 5A, Supplementary Figure 7-8). In LPS-treated male and female rats, the percentage of iNOS^+^ cells showed a consistent elevation trend throughout the cartilage (Supplementary figure 9A), as well as in the central region (male: 13.68%, 95%CI: -23.04 to 50.41%, Hedges’ g = 0.56, *P* = 0.397; female: 12.77%, 95%CI: -4.24 to 29.78%, Hedges’ g = 1.13, *P* = 0.116) (Figure 5B,C). We therefore focused subsequent analysis on the central region to examine the expression patterns of potential mediators. Notably, LPS was not detected in the cartilage layer, suggesting that the observed changes in cartilage were unlikely attributed to direct LPS accumulation in the cartilage. Nevertheless, leptin and OB-R were detected in the condylar chondrocytes of both sexes and treatment groups (Figure 5B,D,E, Supplementary Figures 7-8), confirming the molecular components required for leptin signalling were present within the TMJ cartilage. OB-R^+^ cells were detected in condylar chondrocytes of both sexes, LPS infusion had a significant overall impact on promoting the abundance of OB-R^+^ cells (F(1,12) = 5.79, partial η² = 0.33, P = 0.033), though this effect did not reach significance within either sex individually (Figure 5D). The percentage of leptin^+^ chondrocytes in female LPS groups was elevated by 23.65% (95%CI: 10.72 to 36.58%) compared to female controls (*P* = 0.004), and was also 18.25% (95%CI: -6.79 to 43.30%) higher than that in the male LPS group (*P* = 0.125) (Figure 5E).

**Figure 5.**
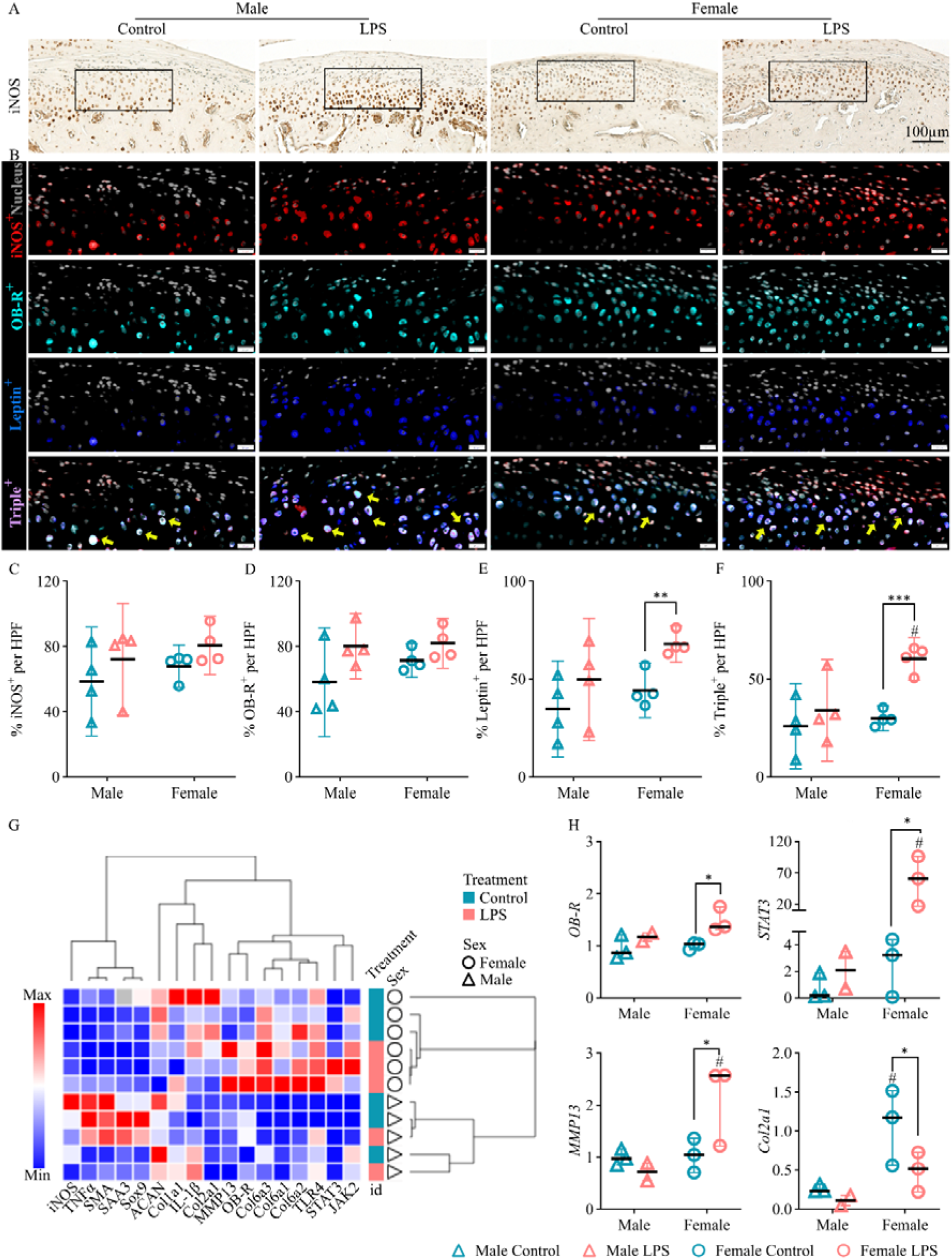
The role of leptin signalling in sex-differentiated OA progression. (A) Representative IHC images showing the expression and distribution of iNOS in cartilage layer. Black box indicating the regions where the high-power field (HPF) of IF images were taken. Scale: 100µm. (B) Representative IF images showing cell nucleus (grey), iNOS^+^ (red), OB-R^+^ (cyan), Leptin^+^ (blue) and Triple^+^ (iNOS^+^OB-R^+^Leptin^+^) cells (pink). Scale: 20µm. Quantitative analysis of the percentage of (C) iNOS^+^, (D) OB-R^+^, (E) Leptin^+^ and (F) Triple^+^ cells over total cells per High Power Field (HPF). N = 4 rats per group and sex. Data represent mean and 95% CI, with each individual data point shown. Comparisons were performed using Student’s t-test. * *P* < 0.05, ** *P* < 0.01 compared to control group within same sex. ^##^ *P* < 0.01 compared to the males within the same treatment group. (G) Heat map depicting gene expression changes of markers associated with LPS, leptin signalling, inflammation, catabolism and cartilage ECM remodelling. The dataset was stratified by sex and clustered according to treatment and genes using one minus Pearson’s correlation for visualization. (H) selected genes that were most influenced by biological sex and LPS. N = 2–3 rats per sex per group. Data are presented as median and range [min – max], with each individual data point shown. Statistical significance was assessed using Two-way ANOVA with Fisher LSD correction. * *P* < 0.05 compared to the control group within the same sex. ^#^ *P* < 0.05 compared to the males within the same treatment group.

OB-R, iNOS and leptin colocalization on chondrocytes appeared in a sex-divergent pattern following LPS infusion. Two-way ANOVA revealed significant main effects of both Female sex (F(1,12) = 7.12, partial η² = 0.37, P = 0.020) and LPS treatment (F(1,12) = 11.53, partial η² = 0.49, P = 0.005) on the percentage of cells stained triple-positive (Triple^+^) for leptin, OB-R and iNOS, with a trending interaction effect (F(1,12) = 3.87, partial η² = 0.24, *P* = 0.073). The % Triple^+^ was significantly higher in LPS-treated females than control females (mean difference: 30.46%, 95%CI: 20.78 to 40.13%, Hedges’ g = 4.74, *P* < 0.001), whereas the increment in males was smaller and statistically non-significant (mean difference: 8.11%, 95%CI: -17.95 to 34.16%, Hedges’ g = 0.47, *P* = 0.48) (Figure 5F). Within the LPS group, female showed 26.33% (95%CI: 4.64 to 48.03%) more Triple^+^ cells than males (Hedges’ g = 1.86, *P* = 0.025). Despite a comparable percentage of iNOS^+^ cells between LPS-treated males and females (HL: 1.13%, *P* = 1.000), 75.75% of iNOS^+^ cells was Triple^+^ in LPS-treated females, as opposed to 52.63% in LPS-treated males (mean difference: 23.12%, 95%CI: - 21.65 to 67.89%, *P* = 0.221) (Supplementary Figure 9B). The differential abundance of Triple^+^ cells between sexes may indicate that in females, iNOS activation was mainly found in chondrocytes co-expressing leptin and OB-R, while in males, nearly half of iNOS^+^ cell population lacked concurrent leptin and OB-R co-expression.

Consistent with immunofluorescence findings showing an increased percentage of triple^+^ cells in LPS-treated females, the expression of genes associated with the leptin receptor (OB-R) and its downstream signalling (STAT3) was significantly upregulated in LPS-treated females compared to same-sex controls. In parallel with changes in leptin signalling, genes related to cartilage matrix remodelling (*MMP13*) were concomitantly upregulated, while *Col2a1* expression was reduced in the female LPS group (Figure 5H and Supplementary Figure 10). Leptin mRNA was not detected in TMJ condylar heads – which comprise cartilage and subchondral bone – from any sex or treatment group, indicating that the leptin protein observed in condylar chondrocytes was unlikely to be locally produced by either tissue but may instead have originated from the systemic circulation.

To further understand the systemic-to-joint link, we pooled samples from the LPS and control groups within each sex for Spearman’s rank correlation analysis, treating leptin concentration as a continuous biological variable to explore its relationships with intra-articular leptin^+^ cells in TMJ cartilage and cartilage degeneration measured by total Mankin score, independent of treatment group assignment. In female rats, circulating leptin was positively correlated with the percentage of leptin^+^ cells in cartilage (rho: +0.76, n = 8, *P* = 0.028), and significantly associated with total Mankin score (rho: +0.77, n = 8, P = 0.024). The intra-articular % Leptin^+^ cells were moderately associated with total Mankin score, although did not reach a statistically significant level (rho: +0.41, n = 8, *P* = 0.319). The findings in female rats support our hypothesis that systemic leptin elevation is associated with greater intra-articular leptin presence and both are related to more severe cartilage damage in females. Discrepancies were noticed in male rats. An inverse association between plasma leptin and intra-articular leptin was observed (rho: -0.31, n = 8, *P* = 0.456), paralleled by negative correlation between plasma leptin and Mankin score (-0.62, n = 8, *P* = 0.105); while intra-articular leptin percentage showed a positive trend in association with Mankin score (rho +0.66, n = 8, *P* = 0.073). The opposing directions in male rats suggest that systemic leptin status does not simultaneously reflect the joint leptin abundance nor cartilage damage (Supplementary Figure 11).

### 3.4. Leptin stimulated sex-divergent metabolic response and oxidative stress in chondrocytes

Next, we performed *in vitro* experiments to investigate the role of leptin and its potential synergistic effect with LPS in female and male chondrocytes. The biological sex of the cells was confirmed by agarose electrophoresis (Figure 6A) prior to assigning them to each experiment (Figure 6B). Using Nile red staining, we observed lipid droplet (LD) accumulation in untreated chondrocytes of both sexes (Figure 6C). Upon treatment with increasing leptin doses, the cells lost their intracellular LDs (Figure 6D). The dose-dependent loss of LDs in female chondrocytes was accompanied by a decline in the metabolic rate measured by MTS assay, with an 8% decrease at 2 ng/ml leptin and a 15% reduction at 20 ng/mL leptin (both *P* < 0.05 compared to untreated female). In contrast, the metabolic rate of male chondrocytes was not affected by changes in LDs, remaining above 95% of untreated male control and significantly higher than that of females at all leptin doses (*P* < 0.05) (Figure 6E). Furthermore, female chondrocytes were more sensitive to leptin-induced oxidative stress than male cells. NO production in female chondrocytes increased by 250% with 5 ng/ml leptin treatment and by over 300% with 10 ng/ml leptin (*P* < 0.05), whereas in male cells, NO production remained at baseline with 5 ng/mL leptin and increased only with 10 ng/mL or higher (Figure 6F).

**Figure 6.**
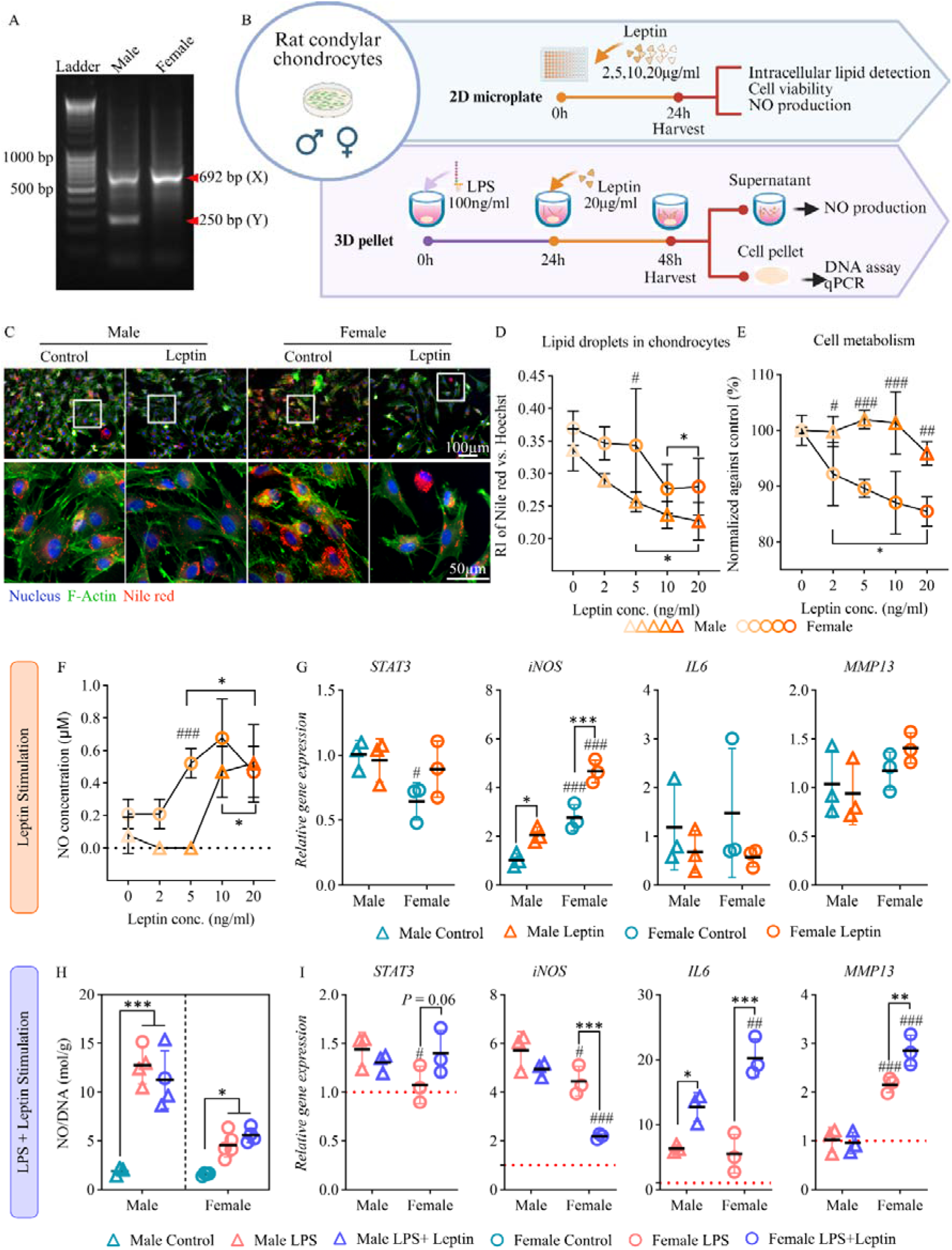
Leptin action in male and female chondrocytes. (A) Agarose electrophoresis to verify the biological sex of the primary rat chondrocytes. (B) Schematic illustration showing the *in vitro* experiment setups. Created in BioRender. Zhang, S. (2026) https://BioRender.com/zbjo4z8. (C) Representative images of Nile red staining showing the changes of lipid droplets in the chondrocytes upon treatment with 20ng/ml of leptin. Lipid droplets were stained with Nile red and shown in red. The cell nucleus was stained with Hoechst and shown as grey. The cell cytoskeleton was stained with F-Actin and shown in green. Top panel: overview of the 2D culture. White boxes indicate where a high magnification image was taken. Scale bar: 100µm. Bottom panel: high magnification images of the chondrocytes. Scale bar: 50µm. N = 3 per group and sex. The dose-dependent changes in (D) intracellular lipid droplet amount, (E) metabolic activity and (F) NO production of chondrocytes treated with varying concentrations of leptin. (G) gene expression of healthy chondrocyte pellets treated with or without 20ng/ml leptin for 24 hours. (H) NO production normalised against the total DNA and (I) relative gene expression normalised against male controls (red dotted line) of the cell pellets primed with or without100 ng/ml LPS for 24 hours, then treated with or without 20ng/ml leptin for 24 hours. Data represent mean ± SD. N = 3 – 5 per group and sex. Statistical significance was assessed Two-way ANOVA with Fisher LSD *post hoc*.* *P* < 0.05, ** *P* < 0.01, *** *P* < 0.001 compared to control group within same sex. ^#^ *P* < 0.05, ^##^ *P* < 0.01, ^###^ *P* < 0.001 compared to males within the same treatment group.

A 3D pellet culture system was used to assess the individual and synergistic effects of leptin and LPS (Figure 6B). Treating healthy cells with 20 ng/ml leptin increased *STAT3* expression in females and significantly upregulated *iNOS* expression in both sexes (Figure 6G). LPS pre-conditioning markedly increased oxidative stress and expression of catabolic genes in chondrocytes in both sexes (Figure 6H). Leptin treatment on LPS-primed cells further increased *IL-6* gene expression in both sexes but only upregulated *MMP13* gene expression in females. Interestingly, leptin also attenuated LPS-induced *iNOS* gene expression in female chondrocytes (Figure 6I).

## 4. Discussion

This study investigated whether chronic systemic endotoxin exposure drives TMJ osteoarthritis development, the involvement of adipose tissue and adipose-derived leptin in this process, and if females and males react differently to the inflammatory/metabolic perturbation. We found that systemic LPS exposure contributed to TMJ arthritic changes and subcutaneous adipose hypertrophy. LPS was absent in cartilage, but intra-articular leptin abundance trended positively with overall TMJ cartilage damage in both sexes. While systemic LPS-triggered joint damage and local response to intra-articular leptin seemed to be shared biological processes with an asymmetric magnitude of response between sexes, the systemic-to-local joint metabolic link was traceable only in female rats, who showed significant elevation of circulating leptin that was largely absent in males. *In vitro*, female chondrocytes were more sensitive to leptin-induced metabolic disturbance. These differences in systemic reactions and chondrocyte responsiveness may, at least partly, explain the sex-divergent pattern of TMJ OA development.

### Systemic LPS infusion associated with different TMJ OA degeneration trajectory in each sex

Gram-negative bacteria-derived LPS is increasingly recognized as a risk factor linking obesity, metabolic dysfunction and knee metabolic OA (20,21,47). Systemic delivery of LPS to rodents has been shown to upregulate adipose inflammatory markers in a manner similar to high-fat diet (15) and exacerbate post-traumatic knee OA without significant sex differences (24). We similarly found that LPS induced mild TMJ cartilage degeneration in both sexes, with total Mankin score differing minimally by sex in LPS-treated animals. However, the LPS-triggered TMJ OA showed different pathological changes between sexes. Under the same LPS insult, females exhibited cartilage GAG loss and collagen fibre derangement, trabecular bone deterioration and synovial inflammation, while male rats showed cartilage surface irregularity without significant changes in bone and synovium. This sex-divergence in TMJ OA subtypes is broadly consistent with clinical observations that female TMJ OA patients developed both cartilage and bone degeneration, while TMJ OA in males mainly affected cartilage without the involvement of bone (4).

### Subcutaneous LPS delivery led to sex-dimorphic systemic response and leptin production

Six weeks of LPS infusion led to an increasing trend in endpoint plasma LPS concentration in LPS-treated female rats compared to female controls, reaching a level comparable to the metabolic endotoxemia (15), while minimal change in plasma LPS was detected in males, similar to a previous mouse study (48). Despite the sex-divergent trend, neither sex showed a statistically meaningful difference in plasma LPS between groups. However, this does not necessarily mean subcutaneously infused-LPS failed to enter blood circulation. Systemic LPS is known to promote systemic low-grade inflammation (47) by triggering the release of pro-inflammatory cytokines (49,50). In this study, we showed that LPS infusion produced an overall trend towards increasing plasma NO, reaching a statistically significant difference in female but not in males, suggesting that LPS exerted a measurable systemic effect despite its own concentration not being significantly elevated at study endpoint.

The pathological effect of LPS was further supported by changes in adipose tissue. We observed significant adipocyte hypertrophy in SAT in both sexes following LPS infusion, consistent with studies showing systemic LPS could directly act on adipose tissue to induce pathological adipose tissue expansion (15,17,51). Despite this shared SAT adipocyte hypertrophic response, the downstream metabolic consequences differ between sexes. Leptin is primarily secreted from white adipose tissue in direct proportion to fat content and correlates more strongly with adiposity in females due to estrogen (52). This sex difference was successfully recapitulated in our study. Circulating leptin increased in females in response to LPS but not in males, in line with clinical report that both adiposity and biological sex were significant determinants of leptin concentration (53,54). The plasma leptin elevation in LPS-infused female rats was accompanied by significant upregulation of *TLR4*, which has been reported to mediate LPS-induced signal transduction (55) and downstream production of TNF-α (56). Pro-inflammatory cytokines, such as TNF-α, were found to mediate adipose tissue inflammation and alter adipokine production, including leptin (57,58). By contrast, male SAT showed no comparable plasma leptin elevation and transcriptional activation.

### Plasma leptin concentration significantly correlated with intra-articular leptin and cartilage alteration in females but not in males

As circulating leptin has increasingly been recognized as a regulator of joint pathophysiology (31,59), we next asked whether the female-biased pattern extended to leptin signalling within the joint. Leptin protein was detectable in condylar chondrocytes in both sexes and treatment groups. Although OA chondrocytes were thought to produce leptin (60), leptin mRNA was undetectable in all condyles in our study, indicating that the leptin found in cartilage most plausibly originates from circulation rather than local production by chondrocytes, though local production by other joint tissues cannot be completely excluded (61). The systemic-to-joint leptin link has been described previously in knee joint (31), and supported by our correlation analysis in females, where circulating leptin concentration positively associated with intra-articular leptin^+^ cells and total Mankin score. In males, however, the relationship between plasma leptin and intra-articular leptin was reversed. The origin of intra-articular leptin in males remains unresolved and requires further investigation. Regardless of the origins, once leptin reached the cartilage, it could bind its receptors (OB-R) on the chondrocytes and trigger NO production (62,63). Consistent with this, we showed a spatial co-localization of leptin signal with OB-R- and iNOS-expressing cells in both sexes, with a significantly greater percentage of triple^+^ cells in LPS-treated females than LPS-treated males and sex-matched controls. Gene expression profiles in TMJ condyles further suggested that leptin might induce the NO production in chondrocytes through OB-R-mediated activation of JAK/STAT3 pathway (62). Nevertheless, LPS was not detected in cartilage at the study endpoint. It remains unclear whether LPS never reached the joint or was cleared from the joint before sampling.

### The sex-divergent effect of leptin on healthy or LPS-primed chondrocytes

*In vitro*, leptin alone induced NO production in chondrocytes of both sexes without LPS presence, mirroring the LPS-independent signalling context inferred *in vivo*. Female chondrocytes were more sensitive to leptin-induced metabolic reduction and oxidative stress than male cells. These differences may be explained by sexual dimorphism in cellular energy metabolism, as previous work suggests that female cells rely more on lipid as an energy source, while male cells prefer glucose as a fuel (64). Consequently, leptin-induced intracellular lipid reduction may impose a greater metabolic burden on female chondrocytes. When applied to LPS-primed chondrocytes, leptin selectively upregulated *MMP13* in females, providing a possible explanation for cartilage matrix loss in female rats (65,66). Interestingly, leptin also significantly dampened LPS-induced *iNOS* gene expression in female chondrocytes, indicating that leptin’s effect on NO production depends on the cell states, and may involve different signalling activations, which remain to be elucidated in future studies. Together, our results implicate the complex sex-dependent role for leptin in healthy and diseased chondrocytes and highlighted the potential involvement of lipid metabolism in this process.

## 5. Limitations/Future Directions

Beyond scientific findings, this study offers a broader methodological contribution. Preclinical research has historically relied disproportionately on male animals or on sex-pooled analyses, a practice now recognized to risk obscuring genuine sex-specific effects (67–69). By analysing males and females separately at every level of this study, from *in vivo* histopathological analysis to *in vitro* cellular responses, we were able to identify a pattern of sex-divergent LPS sensitivity and downstream metabolic response that a combined-sex analysis would likely have missed. However, our study is not without limitation. First, the sample size calculation was based on the plasma LPS concentration, our secondary outcome. Due to sample preparation error, we failed to obtain histology and microCT results from the pilot trial and could not calculate the sample size based on Mankin score, which was our primary outcome. This led to an overly optimistic sample size estimation and in our main trial, comparison between LPS and control group in Mankin score did not reach a statistically significant level in either sex. Despite the small sample size, this study achieved a borderline 77% study power to detect the effect of LPS in Mankin score in female, reflecting a strong catabolic effect, but only 55% power in males, reflecting a weaker and more variable response. Since the deteriorating effect of LPS on cartilage is clearly sex-divergent, a single sample size calculated from a pooled or single-sex effect estimate is unlikely to be appropriate for both sexes. Future studies should therefore estimate the sample size separately for each sex to avoid false negative error or unnecessary use of animals. Beyond the statistical constraints, several aspects of our experiment design could be improved in future studies. Although our data support an association between LPS, leptin signalling and OA pathogenesis, establishing causal relationships will require further *in vivo* studies using leptin-deficient ob/ob mice or leptin receptor-deficient db/db mice or fa/fa rats. Similarly, a comprehensive understanding of LPS- and leptin-associated lipid metabolism in chondrocytes would benefit from dedicated lipidomic profiling (70). Additionally, excessive compressive force is a recognised primary mechanical trigger for TMJ OA that was not incorporated into the present model (13,71). Future studies would be needed to decipher the interaction of LPS and leptin signalling with a mechanically compromised condyle. Lastly, anatomical and biomechanical differences between rat and human TMJs may limit the clinical relevance of our findings, and further validation in larger animal models or in humanised *in vitro* systems will be essential.

## 6. Conclusion

Our findings show that systemic LPS exposure induces sex-divergent TMJ OA changes in rats, with females and males affected through distinct joint tissues and with greater consistency of response in females. This divergence in OA development trajectory may be associated with a leptin-linked inflammatory response spanning systemic adipose tissue and the local joint. *In vitro* experiments further support a sex-differentiated effect of Leptin on healthy and diseased chondrocytes. Overall, our results suggest that leptin, potentially originating from hypertrophic adipose tissue, links systemic endotoxemia to joint damage. This biological cascade engages more clearly and consistently in females than males (Figure 7). This study provides novel insights that may inform future sex-aware and metabolism-targeted therapeutic strategy for TMJ OA.

**Figure 7.**
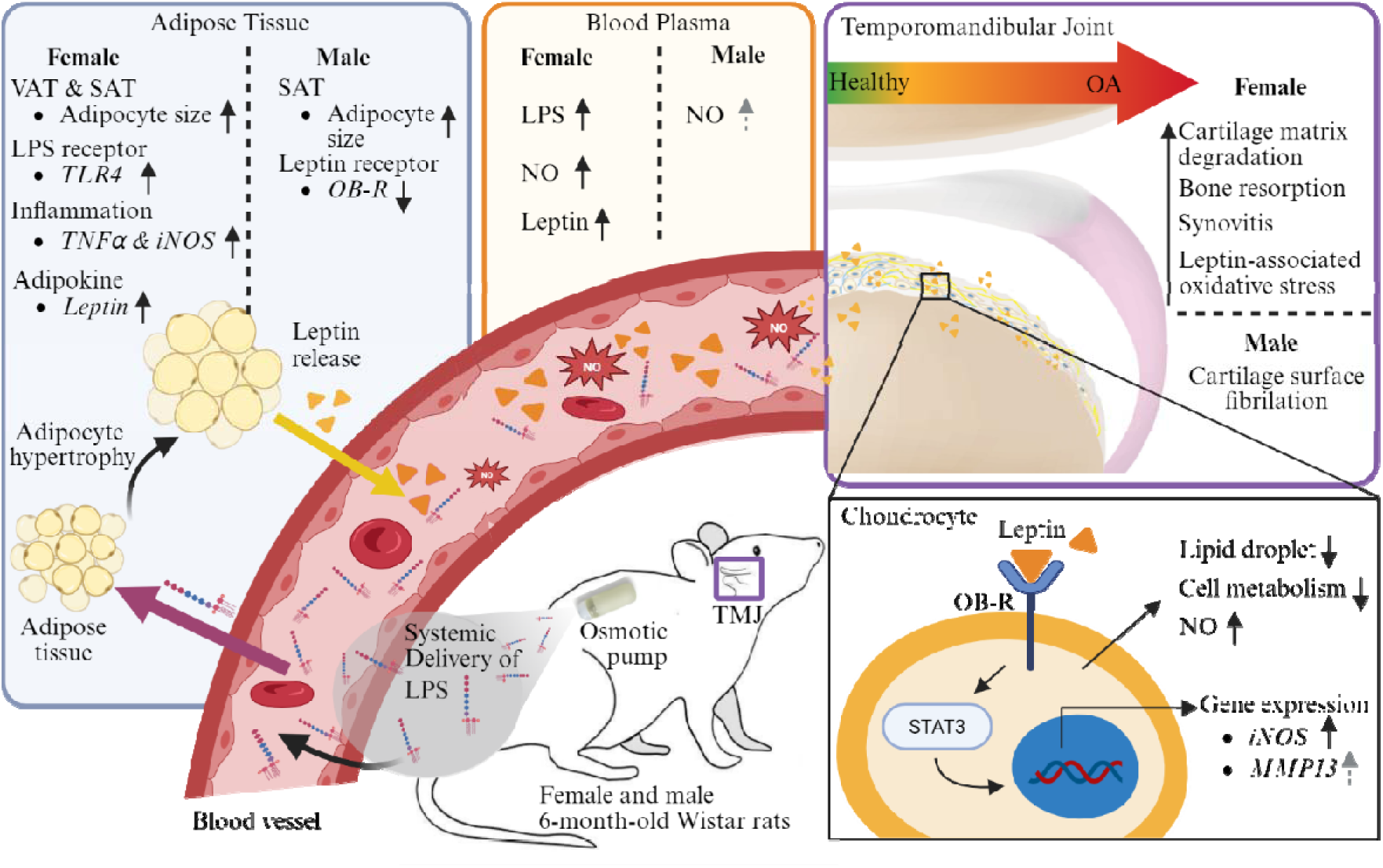
Proposed mechanism of LPS and its interplay with adipose tissue in driving the sex-specific development of TMJ OA. Created in BioRender. Zhang, S. (2026) https://BioRender.com/fttwrav

## 7. Declaration

### Ethics approval and consent to participate

The animal study has been approved by the Veterinary Office of the Canton Zürich (License No. ZH158/2021) and was conducted in accordance with ARRIVE guidelines.

### Consent for publication

Not applicable

### Availability of data and materials

The data that support the findings of this study are openly available in the ETH Zurich Research Collection under the https://doi.org/10.3929/ethz-c-000797498.

### Competing interests

The authors declare that they have no competing interests.

### Role of the Funding Source

This project has received funding from the European Union’s Horizon Europe Research and Innovation Programme under grant agreement No 101095084. This work was supported by the Swiss State Secretariat for Education, Research and Innovation (SERI) under contract number 22.00462.

### Author Contributions

S.Zhang contributed to conception, design, data acquisition, analysis, and interpretation, drafted and critically revised the manuscript; S.Chen contributed to data acquisition, analysis, and critically revised the manuscript; M.Fonti contributed to data acquisition, analysis, and critically revised the manuscript; D.Fercher contributed to design, data acquisition, and critically revised the manuscript. All authors gave final approval and agreed to be accountable for all aspects of the work.

## Acknowledgements

The authors thank the ScopeM facility for technical support with histological sample preparation and microscopy.

## 8. Abbreviation list

ACAN: Aggrecan
ANOVA: Analysis of variance
BMD: Bone mineral density
BMI: Body mass index
BV/TV: Bone volume / total volume
CI: Confidence interval
Col I / Col1a1: Collagen type I / Collagen type I alpha 1
Col II / Col2a1: Collagen type II / Collagen type II alpha 1
Col6a1/Col6a2/Col6a3: Collagen type VI, alpha 1/2/3
DNA: Deoxyribonucleic acid
*E. coli*: *Escherichia coli*
ELISA: Enzyme-linked immunosorbent assay
GAG: Glycosaminoglycan
HE: Haematoxylin and eosin
HFD: High-fat diet
HL: Hodges–Lehmann (location shift estimate)
HPF: High power field
ICC: Interclass correlation coefficient
IF: Immunofluorescence / immunofluorescent
IgG: Immunoglobulin G
IHC: Immunohistochemistry
IL-1β: Interleukin-1 beta
iNOS: Inducible nitric oxide synthase
JAK2: Janus kinase 2
Kdm5c / Kdm5d: Lysine demethylase 5C / 5D
LPS: Lipopolysaccharide
MetOA: Metabolic osteoarthritis
MMP13: Matrix metallopeptidase 13
mRNA: Messenger Ribonucleic acid
MTS: 3-(4,5-dimethylthiazol-2-yl)-5-(3-carboxymethoxyphenyl)-2-(4- sulfophenyl)-2H-tetrazolium (MTS assay)
NaCl: Sodium chloride
NO: Nitric oxide
OA: Osteoarthritis
OB-R: Obesity receptor (Leptin receptor)
PFA: Paraformaldehyde
PSR: Picrosirius red
qRT-PCR: Quantitative reverse transcription polymerase chain reaction
r_rb: Rank-biserial correlation
RNA: Ribonucleic acid
SAA3: Serum amyloid A3
Saf-O: Safranin-O
SAT: Subcutaneous adipose tissue
SD: Standard deviation
SMA: Smooth muscle actin
Sox9: SRY-box transcription factor 9
STAT3: Signal transducer and activator of transcription 3
Tb.N: Trabecular number
Tb.Sp: Trabecular separation
Tb.Th: Trabecular thickness
TLR4: Toll-like receptor 4
TMD: Temporomandibular disorder
TMJ: Temporomandibular joint
TNF-α: Tumour necrosis factor alpha
µCT: Micro-computed tomography
VAT: Visceral adipose tissue
β-Actin: Beta-actin

## Notes

https://doi.org/10.3929/ethz-c-000797498

### Competing Interest Statement

The authors have declared no competing interest.

### Summary of Updates

The manuscript has been revised to update the statistical analysis method, results, and discussion sections. Figures 3 and 4 have been revised.

