## Supplementary file for "Leptin: A Potential Adipokine Bridging Systemic Endotoxemia and Sex-Divergent Temporomandibular Joint Osteoarthritis Development"

**Supplementary Materials**

### 1. Supplementary Tables

Supplementary Table 1. Primer list

| Gene | Primer sequence (5' -> 3') | |
| --- | --- | --- |
|  | Forward | Reverse |
| ß-Actin | GTAGCCATCCAGGCTGTGTT | CCCTCATAGATGGGCACAGT |
| Col1a1 | TCCAGGGCTCCAACGAGA | CTGTAGGTGAATCGACTGTTGC |
| Col2a1 | CCCCTGCAGTACATGCGG | CTCGACGTCATGCTGTCTCAAG |
| Col6a1 | CCCTGGTGGACAAGGTGAAA | CGCATGAGCCCTCTGATGAT |
| Col6a2 | TGGAAGACGTCCTTTGTCCG | GTAGAAGTTCTGCTCGCCCA |
| Col6a3 | GGGACACACGTCTTCAGGTT | CCATGACTGATTGTTGTTGGG |
| TNF-α | CCAGGTTCTCTTCAAGGGACAA | GGTATGAAATGGCAAATCGGCT |
| TLR4 | GCCCTGTTGGATGGAAAAGC | ATGGGTTTTAGGCGCAGAGT |
| JAK2 | GCGACGGGAACAAGATGTGA | TTCAGAACATCGGCCTTCCC |
| STAT3 | TGTGTGACACCATTCATTGATGC | GTTTCCGCTTTCAGCTCCTC |
| MMP13 | CCAGAACTTCCCAACCAT | ACCCTCCATAATGTCATACC |
| ACAN | CAGAACCTTCGCTCCAATGAC | CCTCAATGCCATGCATCACTT |
| OB-R | TTCCTCTTGTGTCCTGCTGC | TCTTGGGGTTTGGAACATCGT |
| SOX9 | CTGAAGGGCTACGACTGGAC | TACTGGTCTGCCAGCTTCCT |
| Leptin | CGGGCTGGAGGATGAACAAA | AGCTGGATGGTGACACAAGG |
| SAA3 | CGCCTACTCGGACATGAGAA | GCATCATAGTTCCCCCGAGC |
| iNOS | CCCAGGAGGAGAGAGATCCG | TGACCTTCCGCATTAGCACA |
| IL-1ß | CCTATGTCTTG CCGTGGAG | CACACACTAGCAGGTCGTCA |
| Kdm5c | TTTGTACGACTAGGCCCCAC | CCGCTGCCAAATTCTTTGG |
| Kdm5d | TGGTGAGATGGCTGATTCC | CCGCTGCCAAATTCTTTGG |

Supplementary Table 2. Plasma LPS concentrations from the pilot trial used for sample size estimation.

| Rat ID | Sex | Group | LPS (EU/ml) |
| --- | --- | --- | --- |
| 236 | Female | Control | 0.29 |
| 235 | Female | Control | 0.31 |
| 230 | Female | Control | 0.24 |
| 238 | Female | LPS | Excluded, hemolysis |
| 234 | Female | LPS | 0.86 |
| 231 | Female | LPS | 0.67 |
| 241 | Male | Control | 0.61 |
| 245 | Male | Control | 0.31 |
| 246 | Male | Control | Excluded, hemolysis |
| 242 | Male | LPS | 0.26 |
| 250 | Male | LPS | 0.20 |

### 2. Supplementary Figures

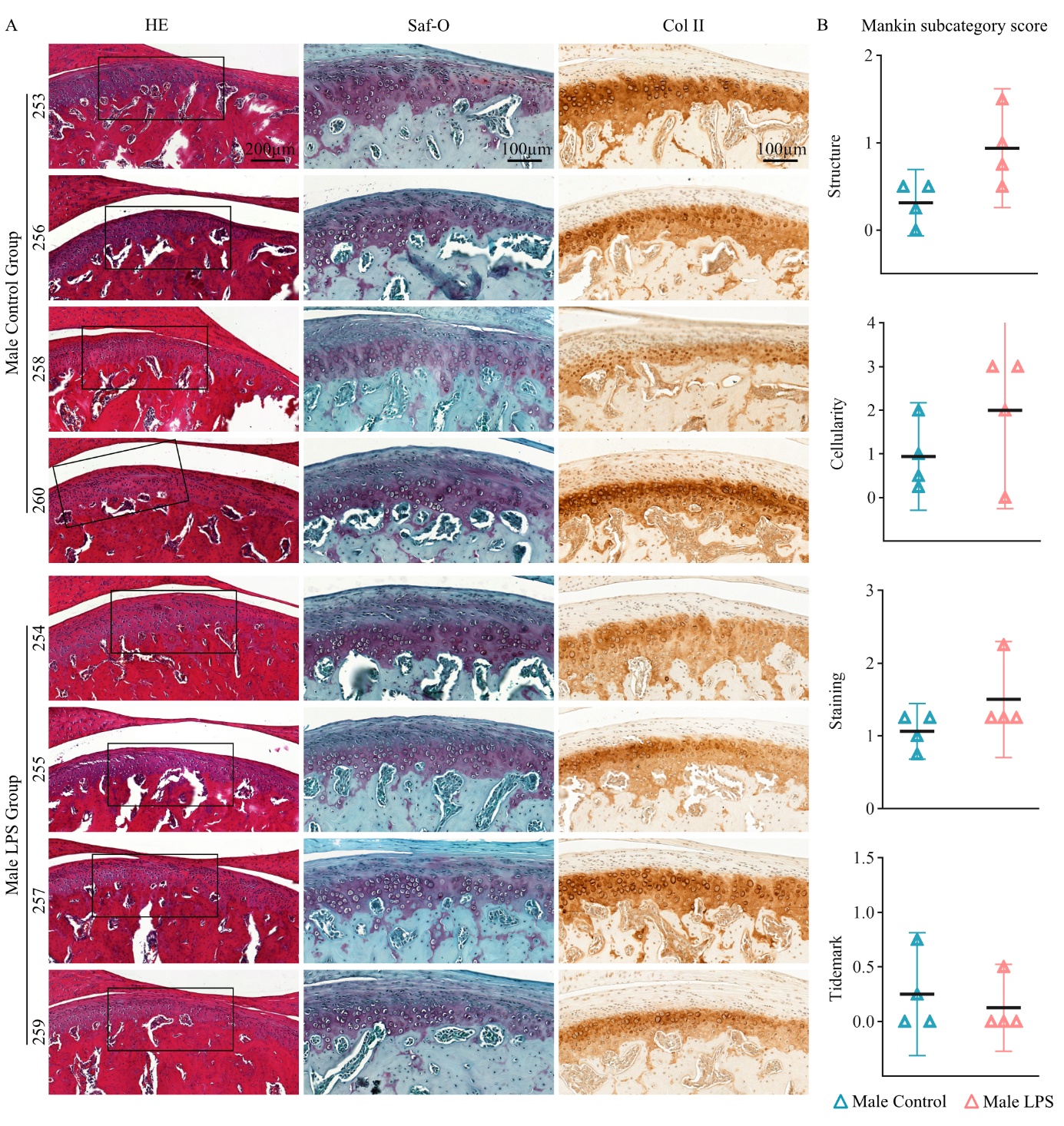

Supplementary Figure 1. (A) Images of TMJ of male rats stained with Hematoxylin and Eosin (HE), Safranin-O and fast green (Saf-O), and collagen II (Col II). Scale bar: 200µm (HE); 100 µm (Saf-O and Col II). (B) Mankin subcategory score for structure, cellularity, staining and tidemark. Data represent the mean with 95% CI. Each data point represents one animal. N = 4 animals per group.

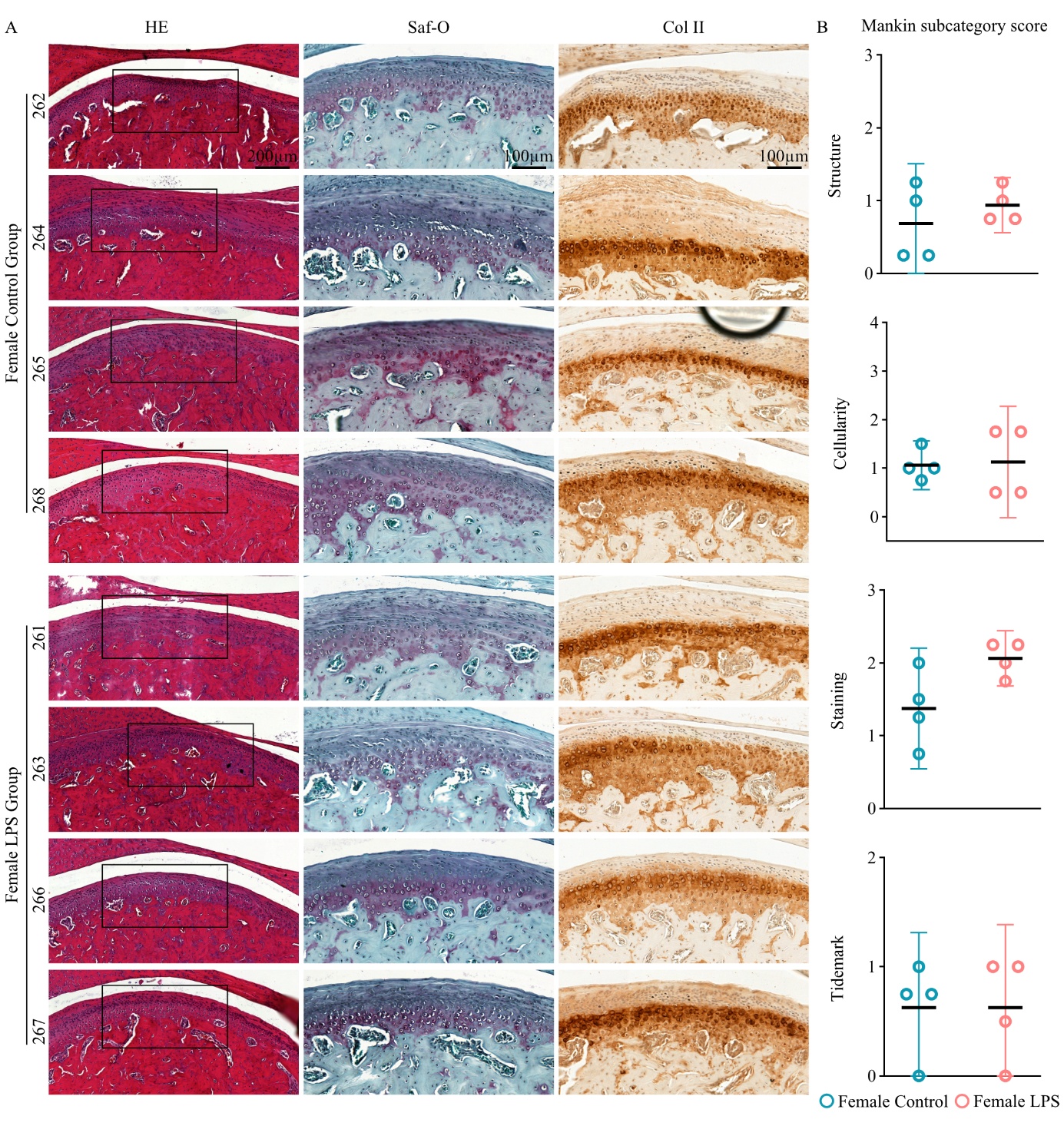

Supplementary Figure 2. (A) Images of TMJ of female rats stained with Hematoxylin and Eosin (HE), Safranin-O and fast green (Saf-O), and collagen II (Col II). Scale bar: 200µm (HE); 100 µm (Saf-O and Col II). (B) Mankin subcategory score for structure, cellularity, staining and tidemark. Data represent the mean with 95% CI. Each data point represents one animal. N = 4 animals per group.

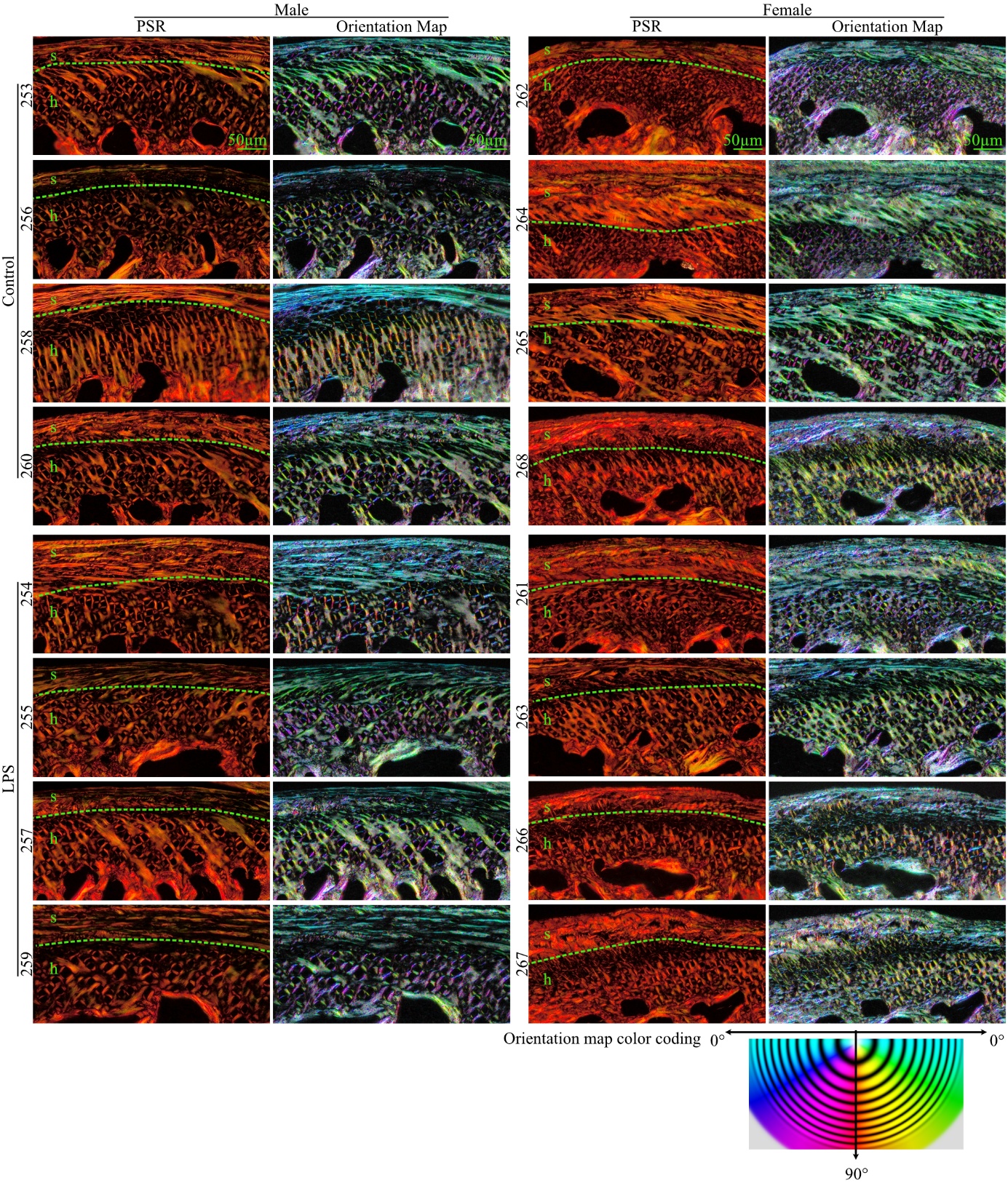

Supplementary Figure 3. Polarised microscopic images of TMJ stained with picrosirius red (PSR), and a pseudo-colour fibre orientation map generated based on PSR images. s: superficial zone; h: hypertrophic zone. Scale bar: 50µm.

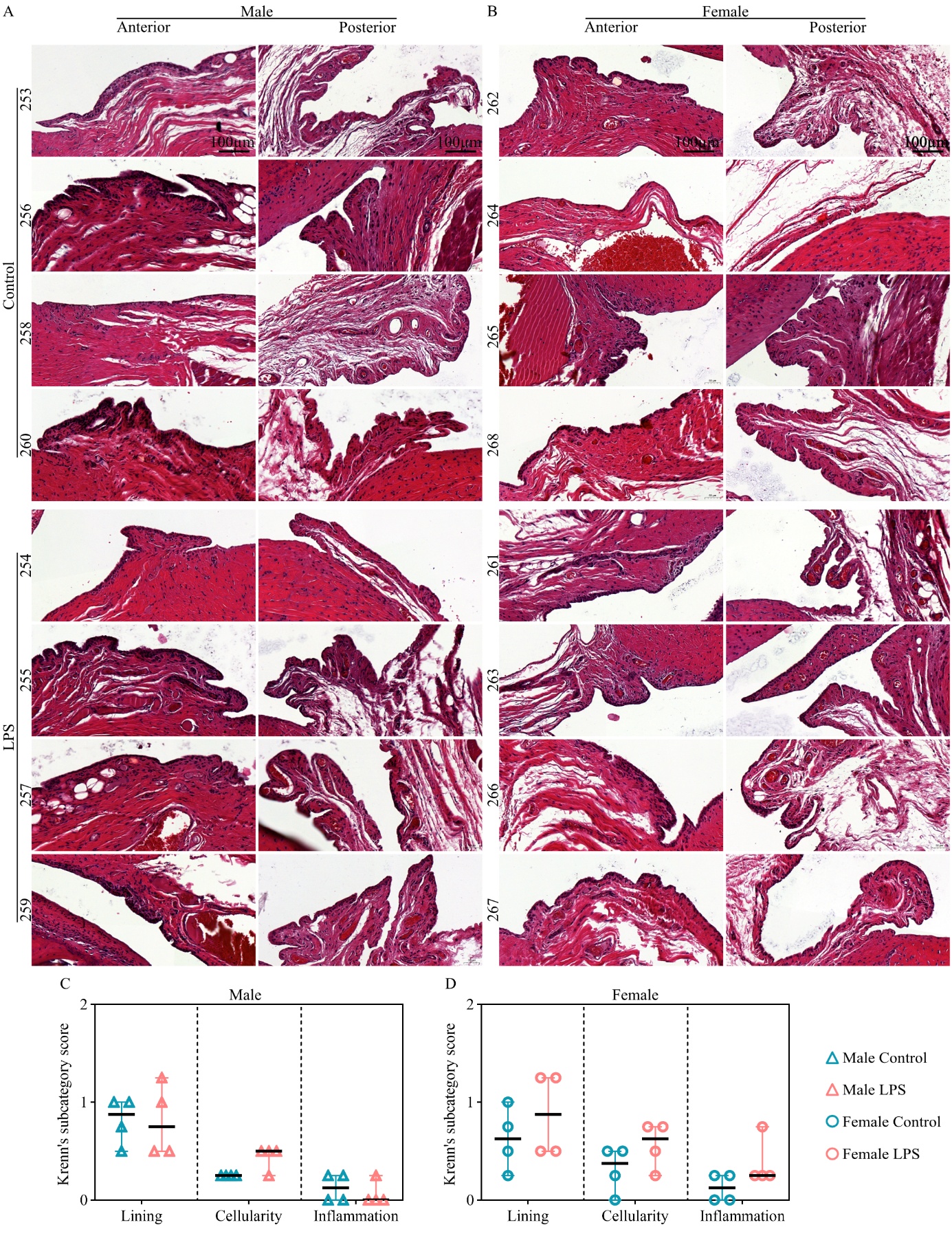

Supplementary Figure 4. (A) Hematoxylin and Eosin (HE) images of anterior and posterior synovial membranes. Scale: 100 µm. (B) Krenn’s subcategory score to evaluate the synovial lining, cellularity, and inflammation. Data represent the mean with 95% CI. Each data point represents one animal. N = 4 animals per group.

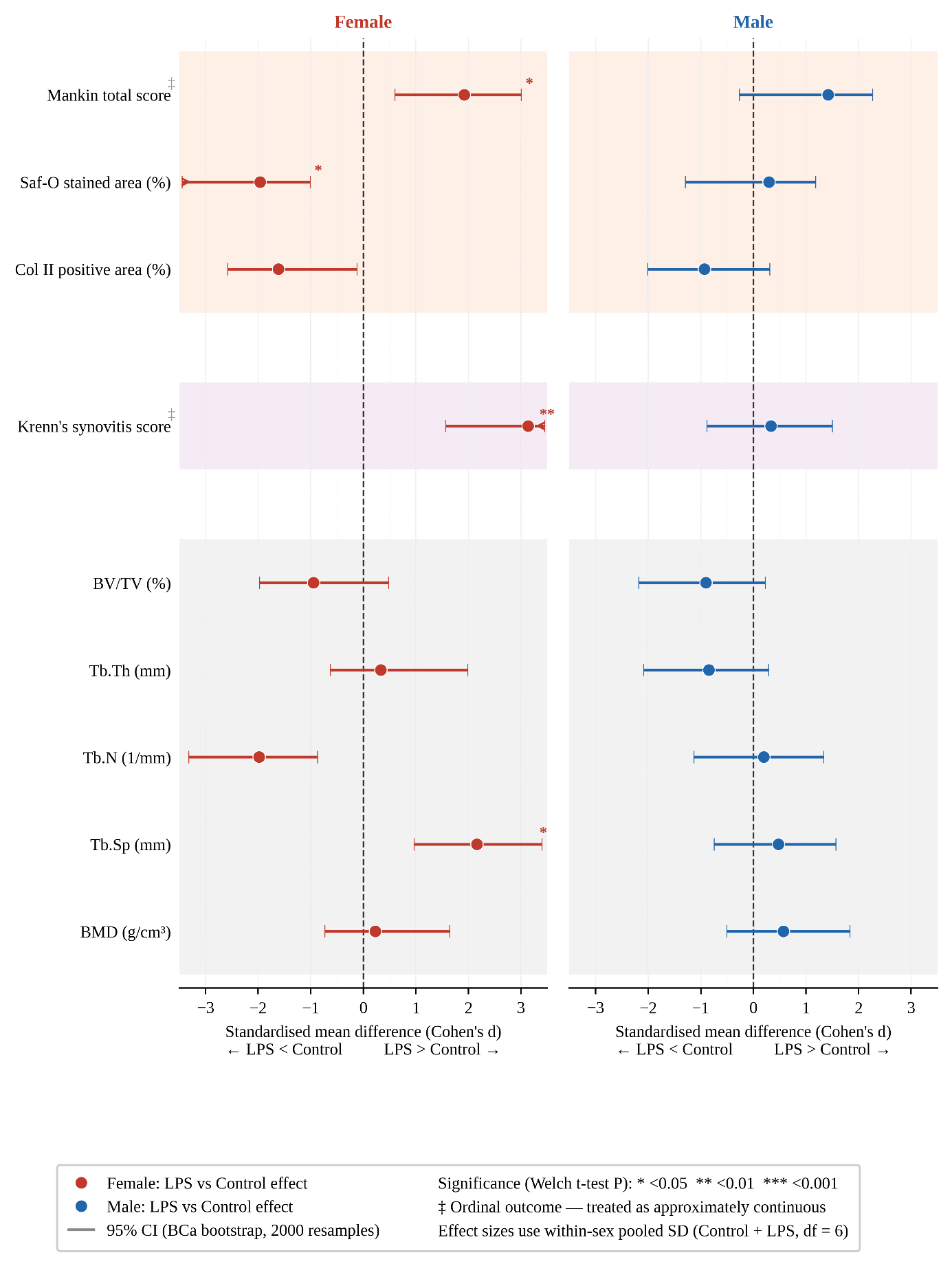

Supplementary Figure 5. Forest plot of standardized effect size for the effect of LPS versus control across joint outcome measures for each sex. Points represent the point estimate of Cohen's d; error bars represent 95% confidence intervals derived from bias-corrected and accelerated (BCa) bootstrap resampling (2,000 resamples). A percentile bootstrap was used for two ordinal outcomes (Mankin score, Krenn's score). Statistical significance was assessed using Welch's t-test (unequal variance). Positive values indicate a higher value in the LPS group relative to control; negative values indicate a lower value. Mankin total score and Krenn's synovitis score (marked ‡) are ordinal outcomes treated as approximately continuous for the purposes of this analysis. *P < 0.05, **P < 0.01, ***P < 0.001.

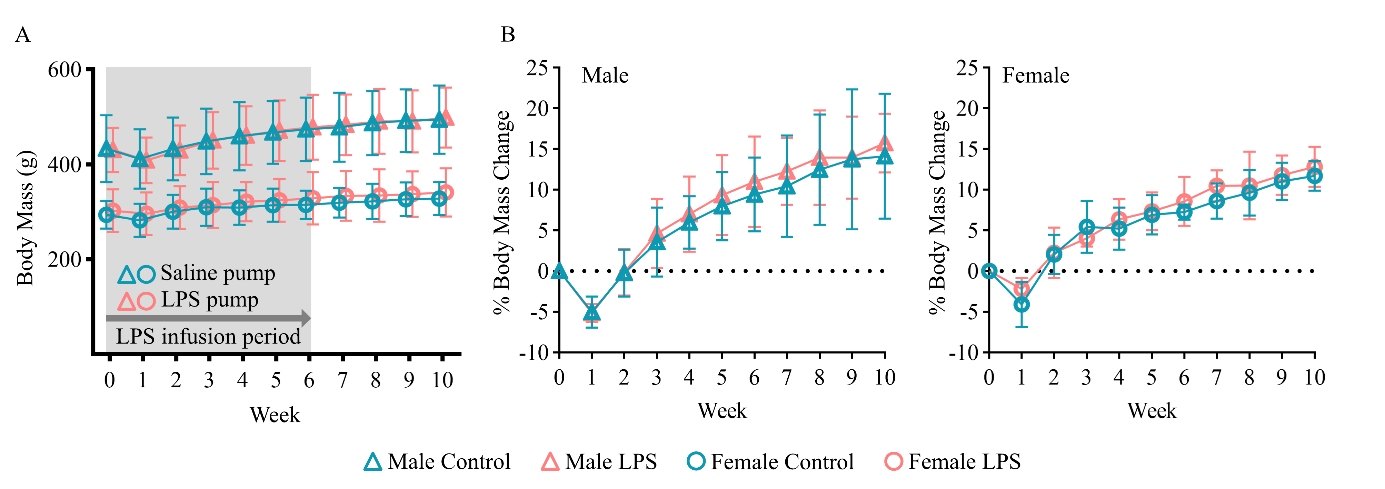

Supplementary Figure 6. Body mass change over time. (A) The absolute body mass of animals and (B) the percentage change in weight compared to week 0. Data represent mean ± 95% CI. N = 4 animals per group and sex.

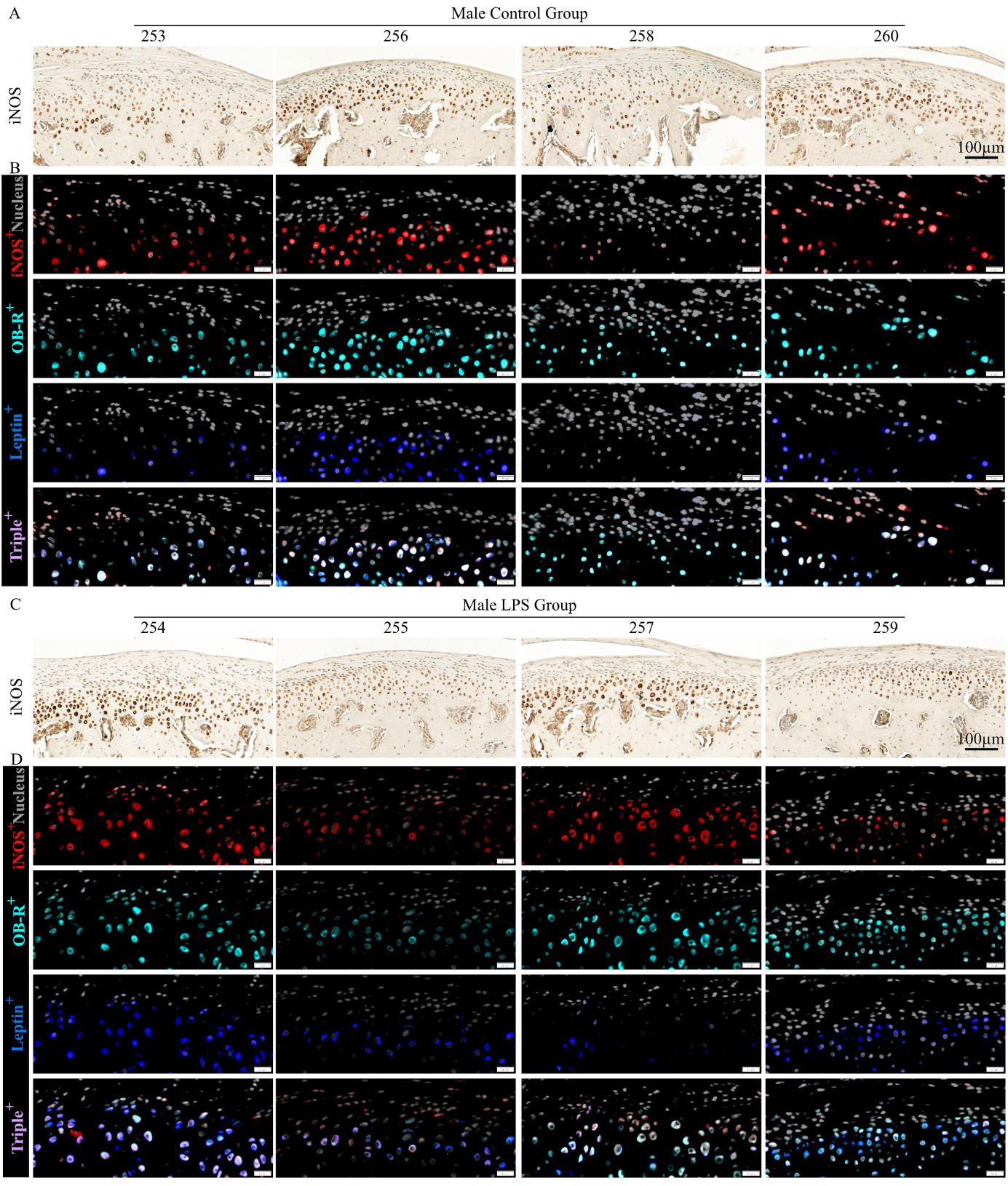

Supplementary Figure 7. The distribution of iNOS, OB-R, and Leptin in male TMJ condylar cartilage. IHC images showing the expression and distribution of iNOS in the cartilage layer of the (A) Control group and (C) LPS group. Scale: 100 µm. Immunofluorescence images of (B) Control group and (D) LPS group showing the cell nucleus (grey), and iNOS^+^ (red), OB-R^+^ (cyan), Leptin^+^ (blue) and triple^+^ (iNOS^+^OB-R^+^Leptin^+^) (pink) cells at central region of the cartilage. Scale: 20 µm.

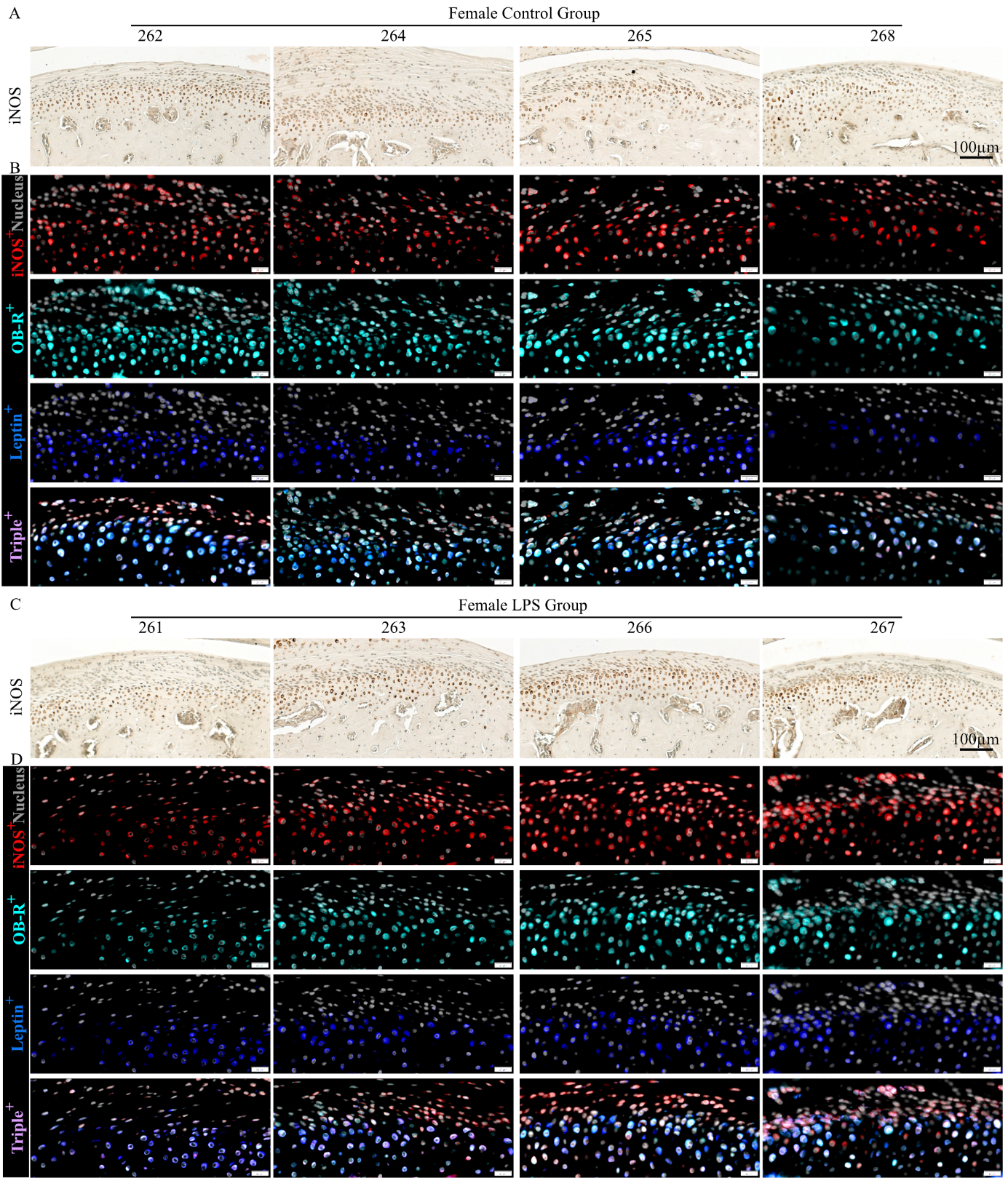

Supplementary Figure 8. The distribution of iNOS, OB-R, and Leptin in female TMJ condylar cartilage. IHC images showing the expression and distribution of iNOS in the cartilage layer of the (A) Control group and (C) LPS group. Scale: 100 µm. Immunofluorescence images of (B) Control group and (D) LPS group showing the cell nucleus (grey), and iNOS^+^ (red), OB-R^+^ (cyan), Leptin^+^ (blue) and triple^+^ (iNOS^+^OB-R^+^Leptin^+^) (pink) cells at central region of the cartilage. Scale: 20 µm.

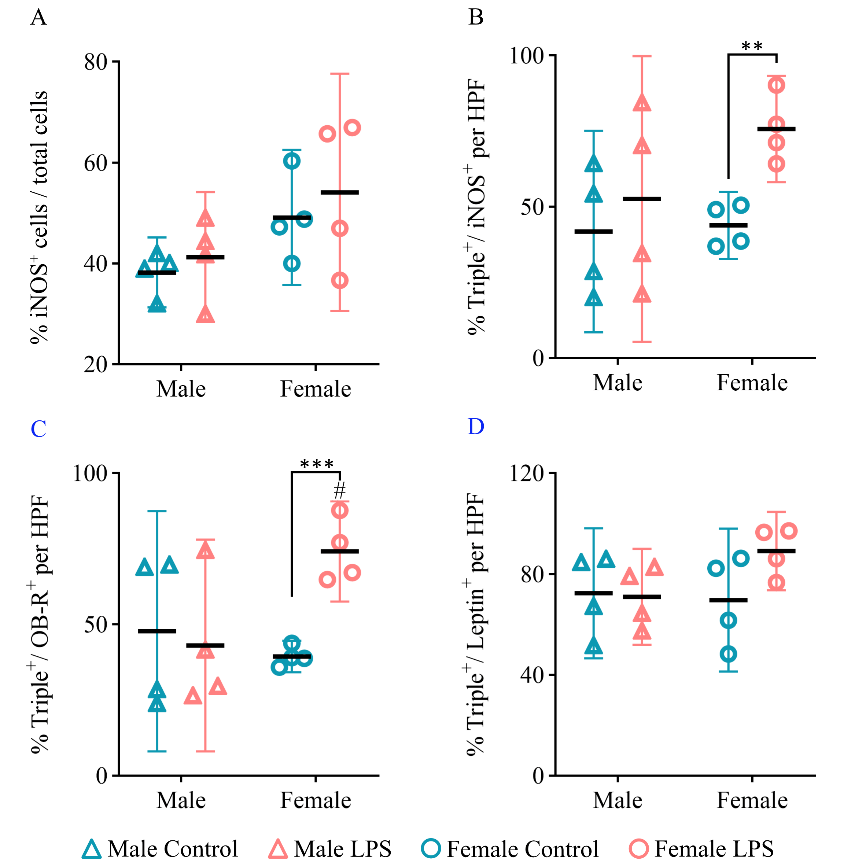

Supplementary Figure 9. Quantitative analysis of the percentage of (A) iNOS^+^ over total cells within the whole condylar cartilage. Percentage of Triple^+^ in subpopulation of (B) iNOS^+^ cells, (C) OB-R^+^ and (D) Leptin^+^ cells per HFP. N = 4 rats per group and sex. Data represent mean with 95% CI. Statistical significance was assessed using student’s t test. ^*^ *P* < 0.05, ^**^ *P* < 0.01 compared to control group within same sex. ^**^ *P*< 0.01, ^***^ *P*< 0.001 compared between LPS and Control groups. ^#^ *P* < 0.05, ^##^ *P* < 0.01 compared to the males within the same treatment group.

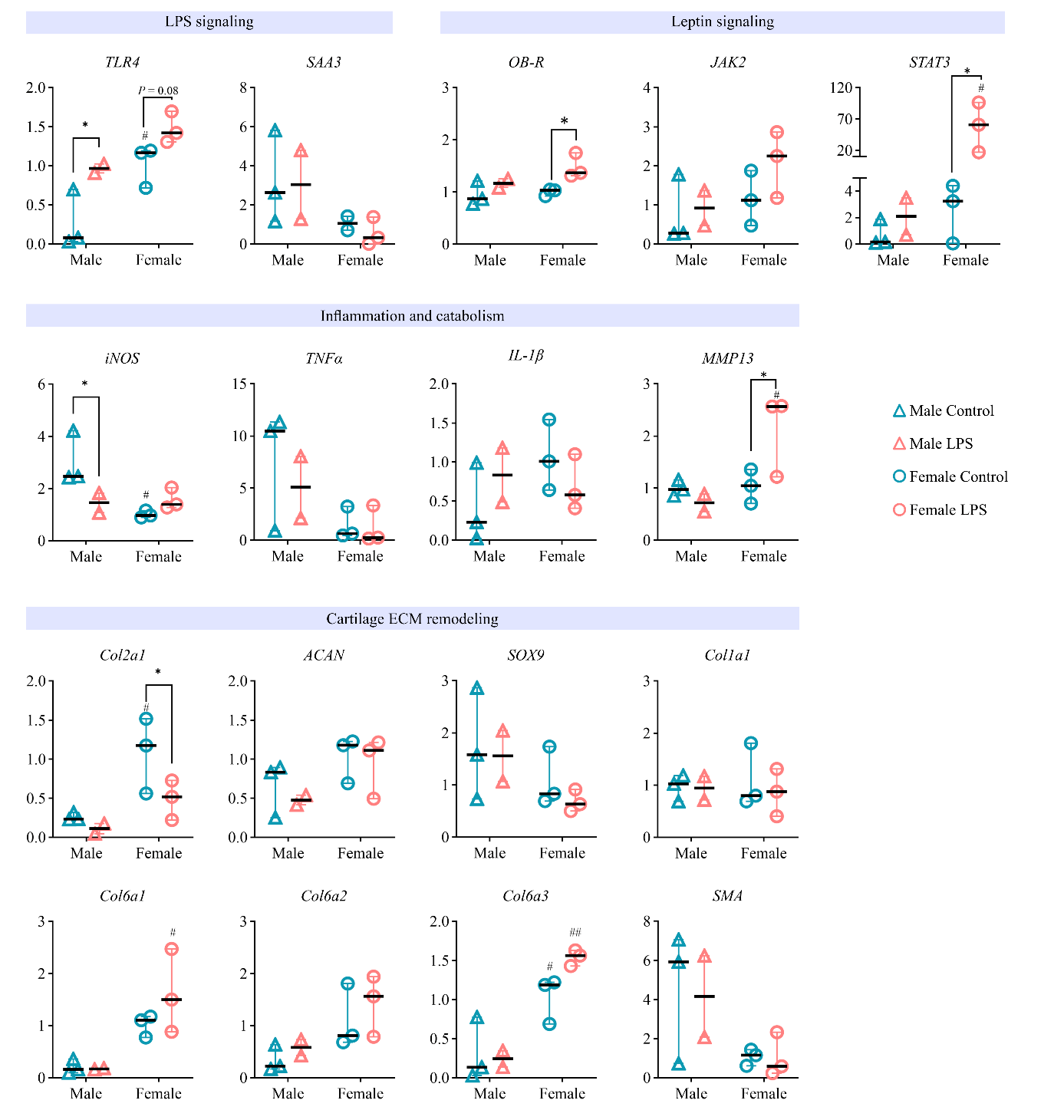

Supplementary Figure 10. Gene expression of TMJ condyle. The relative expression of genes associated with LPS signalling (*TLR4* and *SAA3*), leptin signalling (*OB-R, JAK2* and *STAT3*), inflammation and catabolism (*iNOS*, *TNFα*, *IL-1β* and *MMP13*), and cartilage extracellular matrix (ECM) remodeling (*Col2a1*, *ACAN*, *SOX9*, *Col1a1*, *Col6a1*, *Col6a2* and *SMA*) were compared between sexes and groups. N = 2–3 rats per sex per group. Data are presented as median with range [min-max]. Statistical significance was assessed using Two-Way ANOVA with Fisher LSD *post hoc*. ^*^ *P* < 0.05 compared to the control group within the same sex. ^#^ *P* < 0.05, ^##^ *P* < 0.01 compared to the males within the same treatment group.

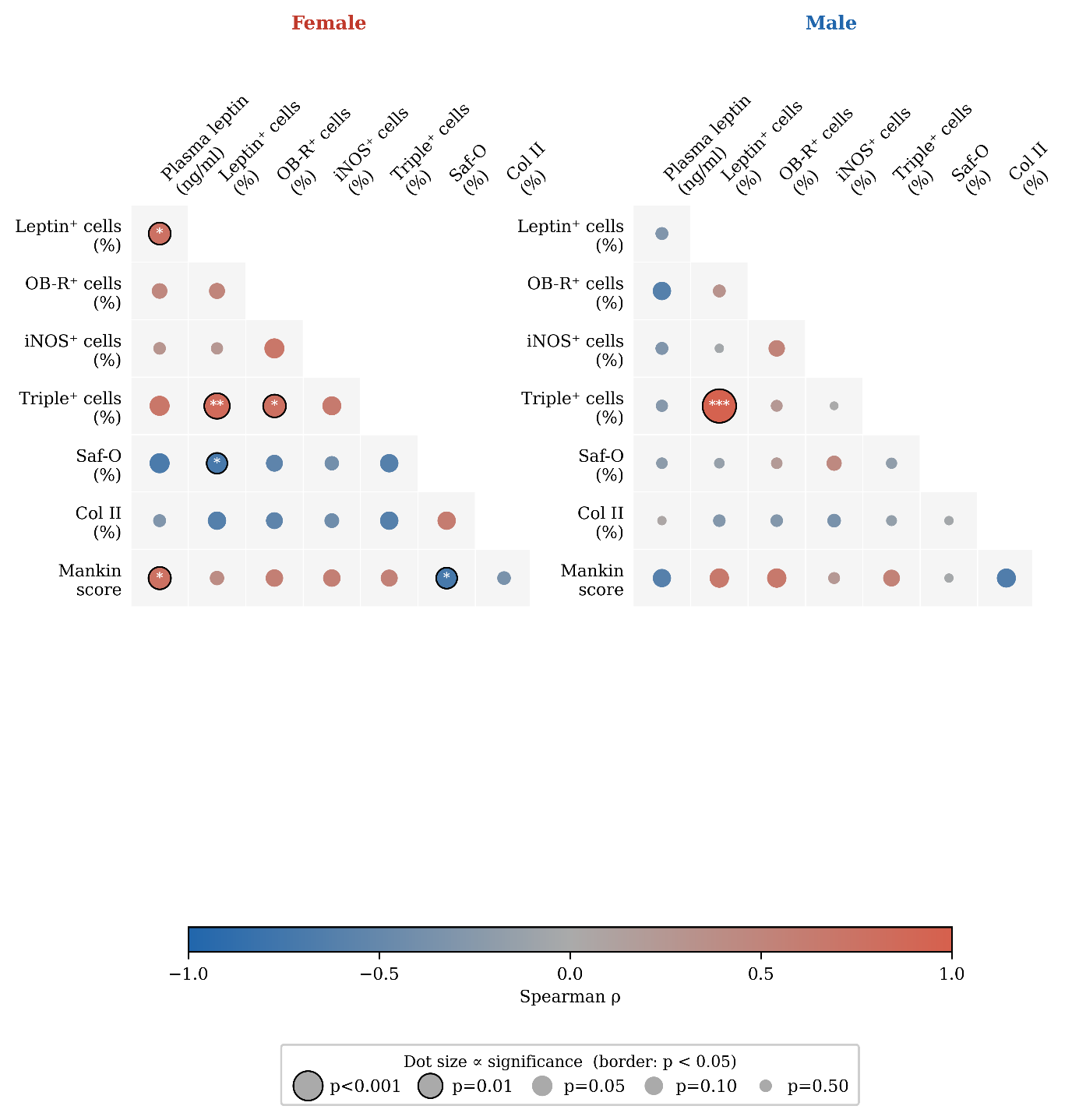

Supplementary Figure 11. Spearman correlation matrix showing the association among systemic leptin concentration, intra-articular leptin^+^ cells (%), and cartilage degeneration measurements in each sex. Pairwise Spearman rank correlations (ρ) were calculated between plasma leptin concentration, the percentage of leptin⁺, OB-R⁺, iNOS⁺, and triple-positive (leptin⁺/OB-R⁺/iNOS⁺) chondrocytes, Safranin-O (Saf-O) stained area, type II collagen (Col II) positive area, and total Mankin score, separately for female and male animals (Control and LPS-treated animals combined within each sex, n = 8/sex). Dot colour indicates the direction and magnitude of the correlation coefficient, from strong negative (blue) to strong positive (red). Dot size is inversely proportional to P-value, such that larger dots indicate stronger statistical significance. A black border denotes *P* < 0.05. ^*^ *P* < 0.05, ^**^ *P* < 0.01, ^***^ *P* < 0.001.

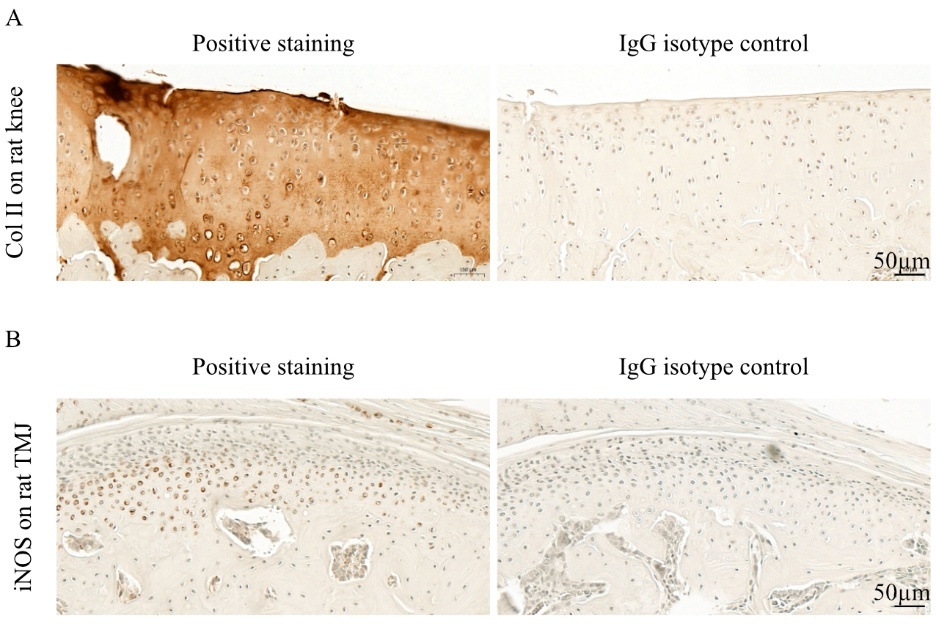

Supplementary Figure 12. (A) Positive staining control and IgG isotype control for Collagen II (Col II) immunohistochemistry. Staining was performed on paraffin sections from the same rat knee. (B) Positive staining control and IgG isotype control for iNOS immunohistochemistry. Staining was performed on paraffin sections from the same rat TMJ. Scale: 50 µm.

### 3. Detailed histological staining and image analysis protocols

White adipose tissue was fixed in 4% PFA for 3 days before histological processing. Following µ-CT examination, the fixed heads were decalcified in 10% aqueous NH4-EDTA for 3 weeks and trimmed to the TMJ region. Then samples were dehydrated to 70 % ethanol, using a series of ethanol solutions with increasing concentrations. After dehydration samples were embedded in paraffin after a Milestone Logos J device (Milestone, ITA) and a Paraffin Embedding Station (Medite TES99). 5 μm-thick sections were cut using a microtome (HM 325, Microm, GER). Sections were then deparaffinized in xylene and rehydrated in several steps from absolute ethanol to pure deionized water before proceeding to the different staining.

#### Hematoxylin and eosin staining

Samples were first stained with Gill’s III hematoxylin for 5 min and washed in running tap water for 10 min. Afterwards, the sections were rinsed in 95% ethanol for 30 s and 40 s in Bluing agent (0.1% Na_2_CO_3_). Finally, they were stained in 0.25% Eosin Y, washed 3 times 1min each in absolute ethanol, dehydrated with xylene, and mounted with Eukitt**^®^** Quick-hardening mounting medium (Merck).

#### Safranin-O staining

First, samples were stained in Weigert’s Iron Hematoxylin solution part A and B 1:1 for 5 min, followed by washing in deionized water and 1% acid alcohol (HCl 37% diluted in 70% ethanol) for 2 s. Then sections were washed again in deionized water, stained in 0.02% Fast Green solution for 1 min, and rinsed with 1% acetic acid for 30 s. Finally, sections were stained in 1% Safranin-O for 30 min, washed in 95% ethanol, dehydrated with xylene, and mounted with Eukitt**^®^** Quick-hardening mounting medium.

#### Picrosirius red staining

The samples were first stained in Weigert’s Iron Hematoxylin solution (part A and B 1:1) for 8 min and washed for 10 min in running tap water. They were then stained in picrosirius red solution for 1 hour, washed in 0.5% v/v acidified water (2.5 ml of acetic acid into 500 ml of deionized water), dehydrated to xylene, and mounted with Eukitt**^®^** Quick-hardening mounting medium.

#### Colorimetric immunohistochemistry (IHC) of collagen II

For Collagen II staining, antigen retrieval was performed with 8000 U/mL hyaluronidase (H3506-1G) in PBS solution for 30 min at 37°C. Sections were subsequently blocked with 5% BSA for 1 hour in at room temperature (RT) and stained with primary antibody II-II6B3 (DSHB Hybridoma Product II-II6B3, 1:20, 75 µg/mL stock) diluted in 1% BSA in PBS overnight at 4°C. The next day, samples were washed twice in PBS for 5 min, then in 0.3% H2O2 for 15 min, twice in PBS for 5 min, and incubated with secondary antibody goat anti-mouse IgG-HRP (ab6789, Goat pAb to Ms IgG (HRP), 1:1000, 2 mg/ml stock) diluted in 1% BSA in PBS for 1 hour. After they were washed 3 times in for 5 min each. Chromogen development was done using DAB substrate kit for 5 min, followed by 3 washing in deionized water for 30 s. Next, nuclei staining was performed with Weigert’s Iron Hematoxylin solution part A and B 1:1 for 5 min, followed by 3 times 30 s wash with deionized water, 2 s in 1% acid alcohol (HCl 37% diluted in 70% ethanol), 3 times 30 s wash with deionized water, 1 min in bluing agent (0.1% Na_2_CO_3_, ), 3 times 30 s wash with deionized water. The slides were then dehydrated in ethanol and xylene and mounted with Eukitt**^®^** Quick-hardening mounting medium. Mouse IgG (Thermo Fisher) was used as a staining control to assess the specificity of the primary antibodies (Supplementary Figure 12A).

#### IHC for iNOS

The antigen retrieval was performed in 10mM sodium citrate buffer at 60°C for 60 minutes, pH 6. Sections were subsequently blocked with 5% BSA for 1 hour at room temperature (RT) and stained with primary anti-iNOS antibody (Abcam, ab15323, 1:100) diluted in 1% BSA overnight at 4°C. The next day, samples were washed twice in PBS for 5 min, then in 0.3% H2O2 for 15 min, twice in PBS for 5 min, then with the secondary antibody (Anti-rabbit 555, A21428, Invitrogen, 1:500) at RT for 1 hour and finally visualised with the DAB chromogen substrate for 2 minutes. The nuclei staining was performed with Weigert’s Iron Hematoxylin as described in section 1.4, and the slides were mounted with Eukitt**^®^** Quick-hardening mounting medium. Rabbit IgG (Thermofisher) was used as a staining control to assess the specificity of primary antibodies (Supplementary Figure 12B).

#### Immunofluorescence staining

Immunofluorescence staining (IF) was performed to detect the expression and colocalization of iNOS, leptin and leptin receptor (OB-R) in TMJ condylar cartilage. Briefly, sections were deparaffinized in xylene and rehydrated in 100%, 95%, 70%, 25% of ethanol and to PBS. Antigen retrieval was performed by cooking the sections at 60 °C for 60 minutes in sodium citrate buffer (pH 6.0). followed by blocking with 5% bovine serum albumin (BSA) for 1 hour. The sections were incubated sequentially with primary antibodies anti-iNOS (1:100), anti-leptin receptor (Invitrogen, MA5-35247, 1:200), and anti-leptin (Thermo Fisher Scientific, bs-0409R, 1:200) at 4°C overnight. After each incubation of primary antibody, the sections were rinsed in PBS with 0.1% (v/v) Tween-20 and stained with either Alexa Fluor 555 goat anti-rabbit IgG (1:500), Alexa Fluor 647 goat anti-rabbit IgG (1:500) or Alexa Fluor 488 goat anti-rabbit (1:500) for 1 hour at ambient temperature. Cell nuclei were counterstained with 4′,6-diamidino-2-phenylindole (DAPI). After the staining, the slides were mounted with aqueous mounting media.

#### Imaging

The HE-, Saf-O-, and IHC-stained sections were scanned using a slide scanner (Panoramic 250; 3DHistech, Hungary) with a 20x objective. Sections stained with picrosirius red were imaged using Slideview TM VS200 (Evident/Olympus, Japan) with 0° rotation angle. The IF-stained slides were imaged using Slideview TM VS200. High-power field images (40x) were taken at the centre of the TMJ condylar cartilage in the slide viewer (OlyVIA 4) as the region of interest for subsequent analysis.

#### Image analysis

QuPath v0 and ImageJ software (National Institutes of Health) were used for all image analysis. On HE-stained adipose tissue, the size of 2000–4000 adipocytes was measured, averaged, and presented as the mean adipocyte size for each sample. The Saf-O- and Col II-stained area in condylar cartilage was measured and normalized against the total area of cartilage layer to calculate the percentage (%) of Saf-O^+^ area and Col II^+^ area, respectively. In IHC-stained sections, the iNOS^+^ cells in the whole cartilage layer were counted and normalized against the total number of cells to calculate the percentage of iNOS^+^ cells, as previously described ^28^. The cartilage fibre orientation in the superficial layer (s), and the hypertrophic layer (h) was examined using the OrientationJ plugin in ImageJ software ^31^. For IF staining, high-power field (HPF) images were taken from the hypertrophic layer at the central region of the cartilage. The individual fluorescence channels were merged in ImageJ. DAPI-stained nuclei, iNOS-positive (iNOS+), Leptin-positive cells (Leptin^+^), OB-R-positive cells (OB-R^+^), and iNOS/Leptin/OB-R triple-positive (Triple^+^) cells were manually counted using the multi-point tool. The percentage of iNOS^+^, Leptin^+^, OB-R^+^, and Triple^+^ cells was calculated relative to the total number of DAPI-stained cells per high-power field.
